# High-speed AFM reveals fluctuations and dimer splitting of the N-terminal domain of GluA2-γ2

**DOI:** 10.1101/2023.12.19.572481

**Authors:** Ayumi Sumino, Takashi Sumikama, Yimeng Zhao, Holger Flechsig, Kenichi Umeda, Noriyuki Kodera, Motoyuki Hattori, Mikihiro Shibata

## Abstract

AMPA glutamate receptors (AMPARs) enable rapid excitatory synaptic transmission by localizing to the postsynaptic density of glutamatergic spines. AMPARs possess large extracellular N-terminal domains (NTDs), which participate in AMPAR clustering at synapses. Nevertheless, the dynamics of NTDs and the molecular mechanism governing their synaptic clustering remain elusive. Here, we employed high-speed atomic force microscopy (HS-AFM) to directly visualize the conformational dynamics of NTDs in the GluA2 subunit with TARP γ2 in lipid environments. HS-AFM videos of GluA2-γ2 in the resting and open states revealed fluctuations in NTD dimers. Conversely, in the desensitized state, the two NTD dimers adopted a separated conformation with less fluctuation. Notably, we visualized individual NTD dimers transitioning into monomers. Furthermore, this NTD-dimer splitting resulted in intersubunit exchange between NTD dimers and an increased number of binding sites with the synaptic protein neuronal pentraxin 1. Therefore, our findings illuminate the significance of NTD dynamics in the synaptic clustering of AMPARs.

## Main

α-Amino-3-hydroxy-5-methyl-4-isoxazole propionic acid (AMPA) glutamate receptors (AMPARs) are a class of ionotropic glutamate receptors (iGluRs) with a crucial role in mediating fast excitatory synaptic neurotransmission in the central nervous system^1–4^. The modulation of AMPAR numbers on the postsynaptic membrane affects basal synaptic activity and synaptic plasticity, ultimately resulting in either long-term potentiation (LTP) or long-term depression (LTD)^5, 6^. Signalling through AMPARs greatly influences synaptic strength, and dysregulation of synaptic signalling by AMPARs has been linked to various neurological disorders^7–9^. AMPARs assemble as tetramers^2, 10, 11^ consisting of subunits GluA1-GluA4 (ref.^12^). GluA1 and GluA2 are the most abundant subunits found in the hippocampus^13, 14^. AMPARs function in tandem with auxiliary subunits, such as the transmembrane AMPAR regulatory protein (TARP) γ2 or stargazin, to regulate the trafficking, localization, clustering, kinetics and pharmacology of the assembled receptor complex^15–17^.

AMPAR displays a tripartite ‘Y’-shaped architecture with a dimer-of-dimers organization that has overall twofold rotational symmetry, as reported by crystallography^11^ and cryogenic electron microscopy (cryo-EM)^18^. The three constituent layers of AMPAR can be distinguished based on their extracellular location: comprising the N-terminal domain (NTD: also called amino-terminal domain (ATD)), the ligand-binding domain (LBD), and the transmembrane domain (TMD)^19–21^ (Fig. 1a). Early single-particle electron microscopy of the native heterotetrameric AMPAR-γ2 complex showed a variety of conformations, mainly reflecting variable separation of the two dimerized NTDs^22^. Moreover, the conformational equilibrium of AMPAR-γ2 was altered by glutamate (Glu), indicating that NTD dimer separation is linked to desensitization^22^, in which the channel closes while Glu remains bound. Recent cryo-EM studies of AMPARs with auxiliary subunits in diverse inhibitor- and ligand-bound states revealed structural alterations in the LBDs and the TMDs in the open and desensitized states^23–27^. These findings led to the proposal of a channel opening mechanism for the ion pore involved in ion permeation^25–27^, thereby improving our understanding of the structural changes in the LBDs and TMDs associated with channel opening. However, cryo-EM structures often exclude the NTD layer due to poor EM density for the NTDs^26–28^.

**Fig. 1:**
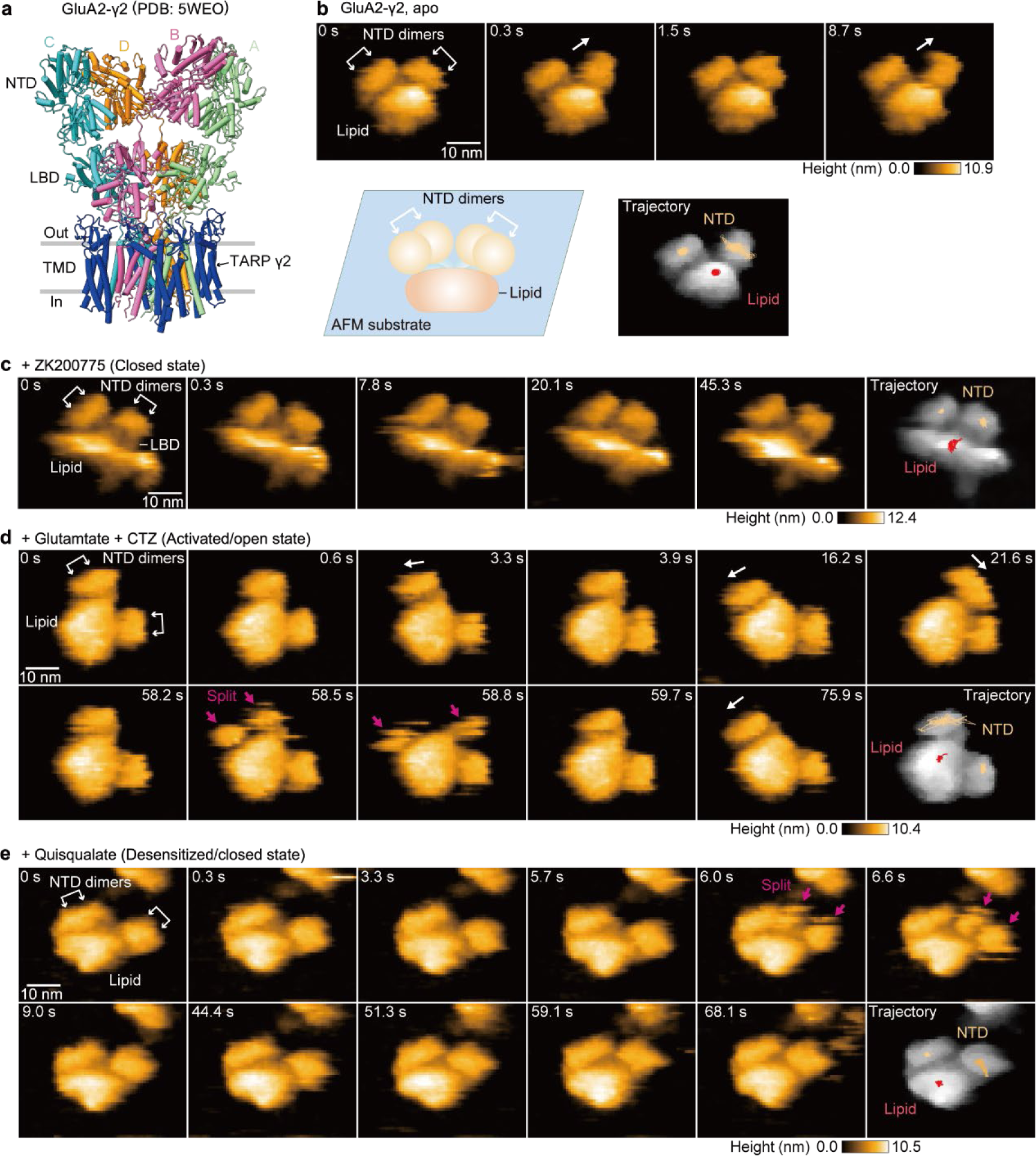
Dynamics of the NTD dimers of GluA2-γ2 visualized by HS-AFM. **a**, Cryo-EM structure of GluA2-γ2 with each subunit in a different colour (PDB: 5WEO). **b-e**, Sequential HS-AFM images of GluA2-γ2 apo (**b**; see also Supplementary Video 1), 0.3 mM ZK200775 (**c**; see also Supplementary Video 2), 3 mM Glu and 0.1 mM CTZ (**d**; see also Supplementary Video 3), and 1 mM Quis (**e**; see also Supplementary Video 4). White arrows indicate the direction of fluctuations of the NTD dimers. Magenta arrows highlight the splitting of the NTD dimers. Trajectories of the NTD dimers and a lipid were tracked for ∼30 s. Imaging parameters: scanning area = 80 × 64 nm^2^ (100 × 80 pixels); frame rate = 3.3 frames/s. The displayed area is 60 × 50 nm^2^. **b**, GluA2-γ2-embedded nanodisc lie down on the mica surface; two NTD dimers are oriented in the same direction relative to the lipid, as shown in a schematic diagram (bottom). White double-headed arrows indicate the NTD dimers. NTD, N-terminal domain; LBD, ligand binding domain; TMD, transmembrane domain; CTZ, cyclothiazide.

Previous studies have reported that NTD-deleted AMPAR mutants still have gating activity but have altered desensitization properties^29, 30^. Furthermore, the NTD exhibits high sequence diversity among GluA1-GluA4 subunits, suggesting the potential for subunit-specific regulation. Additionally, the NTD constitutes roughly half of the AMPAR protein, and its importance has recently been reported, including in subunit assembly^31, 32^, clustering, and localization to synaptic regions^33–37^. The NTD is postulated to serve as a dynamic docking platform within the crowded environment of the synaptic cleft^34^. In particular, neuronal pentraxins (NPs) constitute a family of oligomeric secretory (NP1 and NP2) or membrane-associated (NPR) proteins, which facilitate synaptic clustering via direct interactions between pentraxin (PTX) domains of NP1 (NP1_PTX_) and NTDs^34, 35, 38–40^. However, due to the great structural variability of the NTD, the conformational dynamics of the NTDs remain incompletely characterized and are challenging to observe using ensemble averaging methods such as cryo-EM.

The accumulation of AMPAR structural studies over the past decade has led to proposed activation mechanisms of AMPARs-γ2. However, the conformational dynamics and the structural role of the NTD in synaptic clustering remain unknown. In this study, we employed high-speed atomic force microscopy (HS-AFM) to visualize the conformational dynamics of the NTD of a GluA2-γ2 complex across various functional states, including both the open and desensitized states in lipid environments at room temperature. HS-AFM is a powerful technique capable of observing protein dynamics with nanometre resolution under near-physiological conditions at a timescale of a few hundred millseconds^41–43^. Our HS-AFM videos of GluA2-γ2 revealed fluctuations of the NTD dimers, while these fluctuations were limited in the desensitized/closed state. Additionally, we observed the splitting of the NTD dimers into their constituent monomers. This NTD-dimer splitting caused both intra- and inter-GluA2-γ2 subunit exchange of NTD dimer pairs. Concurrently, an increase in the number of binding sites between the NTD and NP1 was suggested. Our findings indicate that the interreceptor exchange of the dimerized NTDs, combined with the increased number of binding sites to NP1, may serve as a plausible mechanism for promoting the synaptic clustering of AMPARs.

## RESULTS

### NTD fluctuations of GluA2-γ2 in the ligand-free (apo) state

To observe the structural dynamics of AMPARs within the lipid environment, GluA2 was overexpressed in HEK293 cells, purified, and subsequently reconstituted into nanodiscs^44^. HS-AFM observations of GluA2 alone in nanodiscs showed that tetrameric GluA2 collapsed on an AFM substrate. Each GluA2 subunit exhibited two spherical domains, possibly corresponding to the NTDs and LBDs (Extended Data Fig. 1a). To assess their mobility, the radius of gyration (*R*_g_) of the trajectories of the NTDs and the LBDs was calculated, and it was found that the LBDs were less mobile, while the NTDs exhibited extensive movement around the lipid (see Methods and Extended Data Figs. 1b,c). This result implies that the linkers connecting the NTD and the LBD are highly flexible. However, this experimental condition did not accurately depict the dynamics of the GluA2 tetramer due to the large disruption of its steric structure as a channel on the AFM substrate (Extended Data Fig. 1d).

To optimize the experimental conditions, we next expressed and purified a complex between GluA2 and TARP γ2 formed by fusing the N-terminus of TARP γ2 to the C-terminus of GluA2 via a glycine-threonine linker (GluA2-γ2) (Extended Data Fig. 2). Compared to GluA2 alone, the GluA2-γ2 construct exhibits slower deactivation and desensitization, as well as decreased desensitization by electrophysiology, indicating GluA2 modulation by TARP γ2 (ref.^23^). Furthermore, the cryo-EM structure of the GluA2-γ2 construct showed four TARP γ2 molecules bound to the GluA2 tetramer in the mild detergent digitonin^25^, as similarly observed in other GluA2 structures in complex with TARP γ2 in nonfusion constructs^26, 28^. In cryo-EM structures, TARP γ2 interacts with the LBD surface near the lipid, which serves to stabilize the tetrameric structure and regulate the functions of AMPAR^25^. Thus, we hypothesized that this interaction between TARP γ2 and the LBD would facilitate the formation of a stable tetrameric channel structure in GluA2-γ2 even on the AFM substrate. Furthermore, we utilized a fluorochemical surfactant (O310F: (1H, 1H, 2H, 2H-perfluorooctyl)-β-D-maltopyranoside)) because the addition of fluorochemical surfactants at low concentrations around the CMC is often used to improve the orientation bias in single-particle cryo-EM analysis of membrane proteins.

Consequently, HS-AFM images of lipid nanodiscs-reconstituted GluA2-γ2 in the presence of O310F showed two protrusions from the lipid (Fig. 1 and Extended Data Fig. 3). The dimensions of the protrusions were 12.9 ± 1.1 nm horizontally and 9.4 ± 0.8 nm vertically relative to the lipid, with a height of approximately 6.5 nm (Extended Data Figs. 3a,b). Comparison of an AFM simulation and a HS-AFM image of GluA2-γ2 revealed that the dimensions of these protrusions correspond approximately to the size of the NTD dimer of GluA2 (Extended Data Figs. 3c,d). This result indicates that the two protrusions observed in the HS-AFM images originate from the NTD dimers of GluA2-γ2. Furthermore, in some cases, the NTDs formed a structure away from the lipid, and the LBDs were clearly observed between the lipid and the NTDs (Extended Data Fig 3e). The enhanced interaction between TARP γ2 and the LBD is postulated to hinder the dissociation of the tetrameric arrangement on the AFM substrate. Consequently, the GluA2-γ2-embedded nanodiscs did not exhibit discrete unfolding on the AFM substrate; instead, they adsorbed laterally on the AFM substrate (Fig. 1b) and responded differently depending on the ligands applied, presumably reflecting the holding of the physiological nature of the GluA2-γ2-embedded nanodiscs (Figs. 1 and 2). Overall, our utilization of the GluA2-γ2 complex in combination with the application of O310F enabled us to discern the structural dynamics of the NTDs of GluA2-γ2 from a side view at the single-molecule level.

**Fig. 2:**
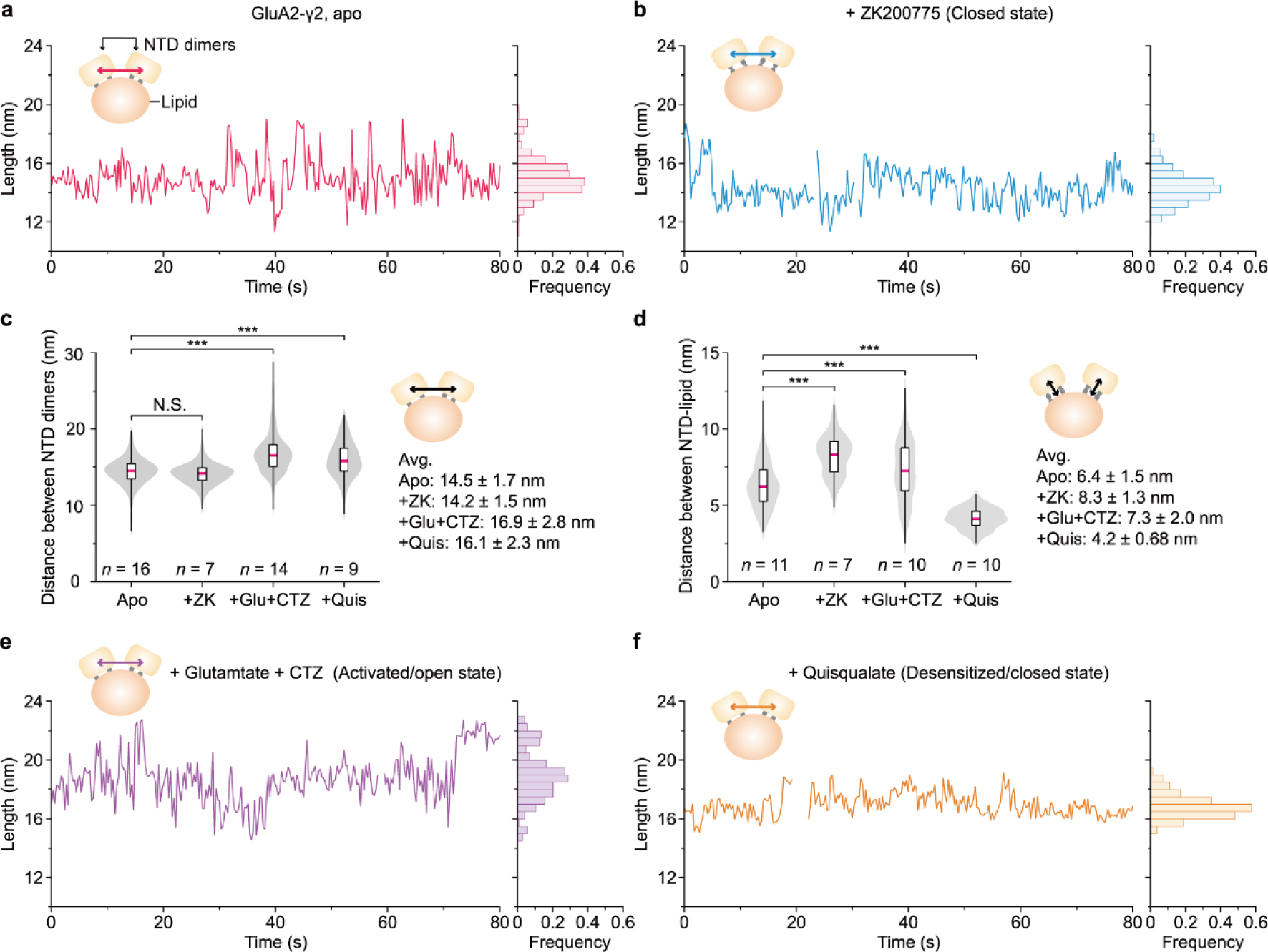
Dynamics of the NTD dimers of GluA2-γ2. **a, b, e, f**, Representative change in distances between the NTD dimers over time in apo GluA2-γ2 (**a**), in GluA2-γ2 with ZK (**b**), in GluA2-γ2 with Glu and CTZ (**e**), and in GluA2-γ2 with Quis (**f**). Blank areas were not measurable due to NTD-dimer splitting. **c**, Distance between the NTD dimers in GluA2-γ2 under different experimental conditions. not significant (N.S.); \*\*\**P* < 0.001 (one-way ANOVA with Dunnett’s post hoc test). **d**, Distance between the NTD dimers and the lipid edge in GluA2-γ2 under different experimental conditions. \*\*\**P* < 0.001 (Kruskal‒Wallis test with Dunnett’s post hoc test). HS-AFM experiments were repeated at least three times independently with similar results. ZK, ZK200775; Glu, glutamate; CTZ, cyclothiazide; Quis, quisqualate.

HS-AFM videos of apo-GluA2-γ2 showed the horizontal fluctuation of one of the two NTD dimers with respect to the lipid edge (Figs. 1b and 2a, Extended Data Fig. 4a, and Supplementary Video 1). In previous structural analyses of GluA2, ZK200775 (ZK), a competitive antagonist with high affinity^45^, has been widely employed to capture the closed formation^11, 18, 23^. Therefore, to capture the structural dynamics of closed GluA2-γ2 under nearly physiological conditions, we performed HS-AFM observations of GluA2-γ2 in the presence of 0.3 mM ZK (GluA2-γ2_ZK_). HS-AFM videos of GluA2-γ2_ZK_ showed similar fluctuations in the NTD dimers as observed in the apo state (Figs. 1c and 2b, Extended Data Figs. 4b and 5, and Supplementary Video 2). Furthermore, although the interdimer distance between the NTDs was comparable to that of the apo state (Fig. 2c), the two NTD dimers were positioned approximately 1.9 nm farther from the lipid edge than in the apo state (apo: 6.4 ± 1.5 nm, ZK: 8.3 ± 1.3 nm in Fig. 2d). Consequently, the LBDs were clearly discernible (Fig. 1c). A previous cryo-EM study demonstrated an extended conformational change between the NTD and the LBD after ZK binding^25^. This consistency with the HS-AFM results confirms the correctness of our measurements (Fig. 3).

**Fig. 3:**
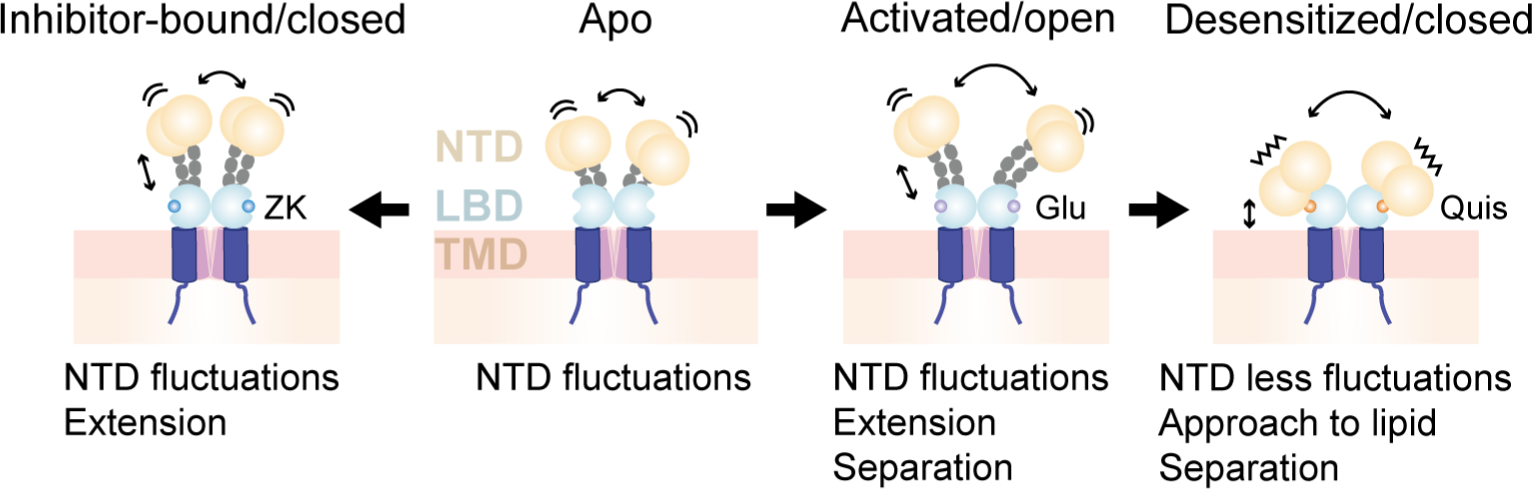
Schematic of the conformational dynamics of the NTD dimers in different ligand-binding states as observed by HS-AFM. The inhibitor-bound/closed, apo, activated/open, and desensitized/closed states are the results from HS-AFM of GluA2-γ2_zk_, GluA2-γ2, GluA2-γ2_Glu+CTZ_, and GluA2-γ2_Quis_, respectively. In the apo state, the NTD dimers exhibit fluctuations parallel to the lipids. Similarly, in the inhibitor-bound/closed state, the NTD dimers also exhibit fluctuations and extended from the lipids. In the activated/open state, the NTD dimers continue to exhibit parallel fluctuations with large separations. In the desensitized state, the NTD dimers show less fluctuation, tending to approach the lipids and separate from each other. ZK, ZK200775; Glu, glutamate; Quis, quisqualate.

### Expanded spatial separation between the NTD dimers of GluA2-γ2 in the activated/open state

Next, to observe the structural dynamics in the activated/open-state GluA2-γ2 (GluA2-γ2_Glu+CTZ_), we added glutamate (Glu), an endogenous agonist for AMPARs, together with cyclothiazide (CTZ)^46^, a positive allosteric modulator of AMPARs, and performed HS-AFM observations. Both Glu and CTZ were present in the imaging buffer under these experimental conditions. This enabled us to capture AMPAR in the open state by inhibiting the transition to the desensitized state^21^. HS-AFM videos of GluA2-γ2_Glu+CTZ_ revealed a notable configuration in which the two NTD dimers adopted a widely separated conformation (Figs. 1d and 2e, Extended Data Fig. 6a, and Supplementary Video 3). One of the two NTD dimers exhibited horizontal fluctuations towards the lipid edge, similar to those in the apo state (Extended Data Fig. 5), and the distance between the two NTD dimers increased significantly by approximately 2.4 nm compared to that in the apo state (apo: 14.5 ± 1.7 nm, +Glu+CTZ: 16.9 ± 2.7 nm in Fig. 2c). Moreover, the distance between the NTD dimers and the lipid edge significantly increased by approximately 0.9 nm in comparison to that in the apo state (apo: 6.4 ± 1.5 nm, +Glu+CTZ: 7.3 ± 2.0 nm in Fig. 2d).

Recent cryo-EM analyses of the GluA2-γ2 complex have revealed structural changes in channel gating, particularly in the structures of the LBD and the TMD in the activated/open state^26, 27^. However, to improve the resolution of cryo-EM structures by class averaging, the NTD layer has typically been excluded due to its heterogeneity and conformational mobility. There is only one structural analysis of the GluA2-γ2 complex in the activated/open state where the NTD layer was not excluded^25^ (PDB: 5WEO or 6DLZ), but the two NTD dimers in the GluA2 tetramer seemed to maintain their relative relationship, and the distance between the two NTD dimers and the edge of the TMD is also essentially maintained. Thus, our HS-AFM results provided unique dynamic features of the NTDs in channel activation, including the large separation of NTD dimers and their disparate distribution relative to the lipid layer (Fig. 3).

### The two NTD dimers of GluA2-γ2 in the desensitized/closed-state exhibited large separation, close to the lipid edge, and adopted a rigid conformation

To visualize desensitized/closed-state GluA2-γ2, we next performed HS-AFM observations of GluA2-γ2 in the presence of L-quisqualate (Quis), a full agonist with high binding affinity. In HS-AFM videos of GluA2-γ2 with Quis (GluA2-γ2_Quis_), the spatial separation between the two NTD dimers was significantly larger than that in the apo state (Fig. 1e and Supplementary Video 4), with an increase of approximately 1.6 nm (+Quis; 16.1 ± 2.3 nm in Fig. 2c). This separation closely resembled the distance observed in GluA2-γ2_Glu+CTZ_. Furthermore, the distance between the NTD dimers and the lipid edge showed a significant decrease of approximately 2.2 nm compared to the apo state (+Quis; 4.2 ± 0.7 nm in Fig. 2d). In contrast, we observed limited horizontal fluctuations of the NTD dimers relative to the lipid edge in GluA2-γ2_Quis_ (Fig. 2f and Extended Data Figs. 5 and 6b). These findings suggest that the NTD dimers of GluA2-γ2_Quis_ assume a rigid conformation with reduced fluctuations as they approach the lipid edge and undergo spatial separation. This lipid-approaching separated conformation resembles the findings from an earlier single-particle EM study of native AMPARs with the TARP γ2 complex^22^. Moreover, the recent cryo-EM analysis of GluA2-γ5 in the desensitized state revealed the NTD configuration when approaching the lipid^47^. Notably, in addition to the consistency of the static architecture, our HS-AFM analysis provided insights into the restricted mobility and rigid conformation of the NTD dimers in the desensitized state (Fig. 3).

### The NTD dimers occasionally dissociate into monomers

Remarkably, our HS-AFM observations revealed the splitting of the NTD dimers into monomers as part of their structural dynamics (58.5 s, 58.8 s in Fig. 1d, and 6.0 s, 6.6 s in Figs. 1e and 4a). This NTD-dimer splitting was consistently observed across all GluA2-γ2 states (Fig. 4b and Extended Data Fig. 7). To quantify the lifetime of the NTD dimer states (stable or split) under different experimental conditions, we computed time-course correlation functions for the transitions between stable and split states of the NTD dimers.

**Fig. 4:**
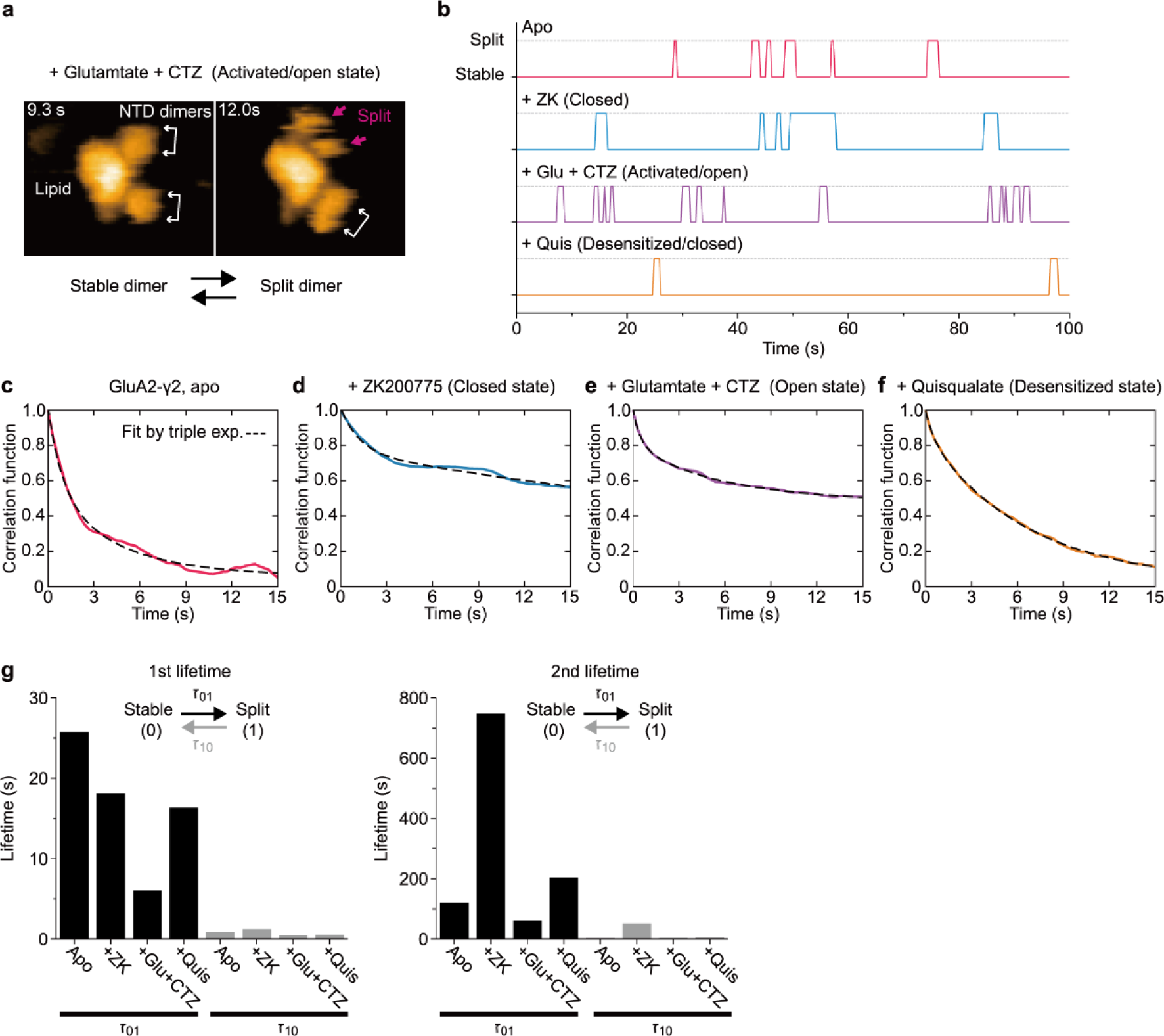
Kinetics of NTD-dimer splitting of GluA2-γ2. **a**, Representative HS-AFM images of NTD-dimer splitting. **b**, Representative time courses of transition of stable and split-states of the NTD dimers under different experimental conditions (see also Extended Data Fig. 7). Dotted lines indicate the split state. **c-f**, Correlation function of NTD-dimer splitting in the apo form (**c**), with ZK (**d**), with Glu+CTZ (**e**), and with Quis (**f**). Each correlation function was fitted by a triple exponential curve, as indicated by the dashed lines. **g**, The 1^st^ (left) and 2^nd^ (right) lifetimes of stable and split-states of the NTD dimers under different conditions (see also Extended Data Table 1). ZK, ZK200775; Glu, glutamate; CTZ, cyclothiazide; Quis, quisqualate.

Lifetimes were calculated by fitting the correlation functions with exponential functions (see Methods and Figs. 4c-f). Our findings showed that the lifetime of the stable state of the NTD dimers in GluA2-γ2_Glu+CTZ_ was approximately more than twice as short as that of the apo state (τ_01_ in Fig. 4g and Supplementary Table S1). Conversely, the lifetime of the split-state of the NTD dimers remained comparable across the three experimental conditions (t_10_ of apo, +Glu+CTZ, and Quis, in Fig. 4g). These results indicate that the NTD dimers of the activated/open state in GluA2-γ2 tend to undergo a transition into the split state (i.e., monomer).

Based on the previous biochemical analysis^32, 48^ and structural study^19^, it seems that the NTD typically forms a stable dimer with extensive interactions. However, it has also been noted that NTDs can be highly dynamic^32, 49–51^, and the possibility of NTD-dimer splitting of AMPARs at room temperature in solution is controversial. To evaluate the energy barriers to this potential dissociation, we computed a free energy profile against the monomer–monomer distance (reaction coordinate, RC) using molecular dynamics (MD) simulations with umbrella sampling (see Methods, Fig. 5, Supplementary Videos 5, 6 and Fig. S1 and S2). The resulting computed free energy profile exhibited two stable positions. The first position, situated at RC = 3.6 nm, aligns with the cryo-EM structure (Figs. 5a,b). We adjusted it so that the structure at RC = 7.0 nm corresponded to the reference energy level (Figs. 5c,d). The second position is located at RC = 9.2 nm, with a free energy well of −1.5 kcal/mol, where the monomers achieve complete separation (Fig. 5f). Notably, in this configuration, one monomer remains in interaction with the undivided dimer, while the other monomer is oriented towards the lipid and occasionally forms interdimer subunit exchange described later. These results indicate that the NTD dimers possess a metastable structure even in the split state.

**Fig. 5:**
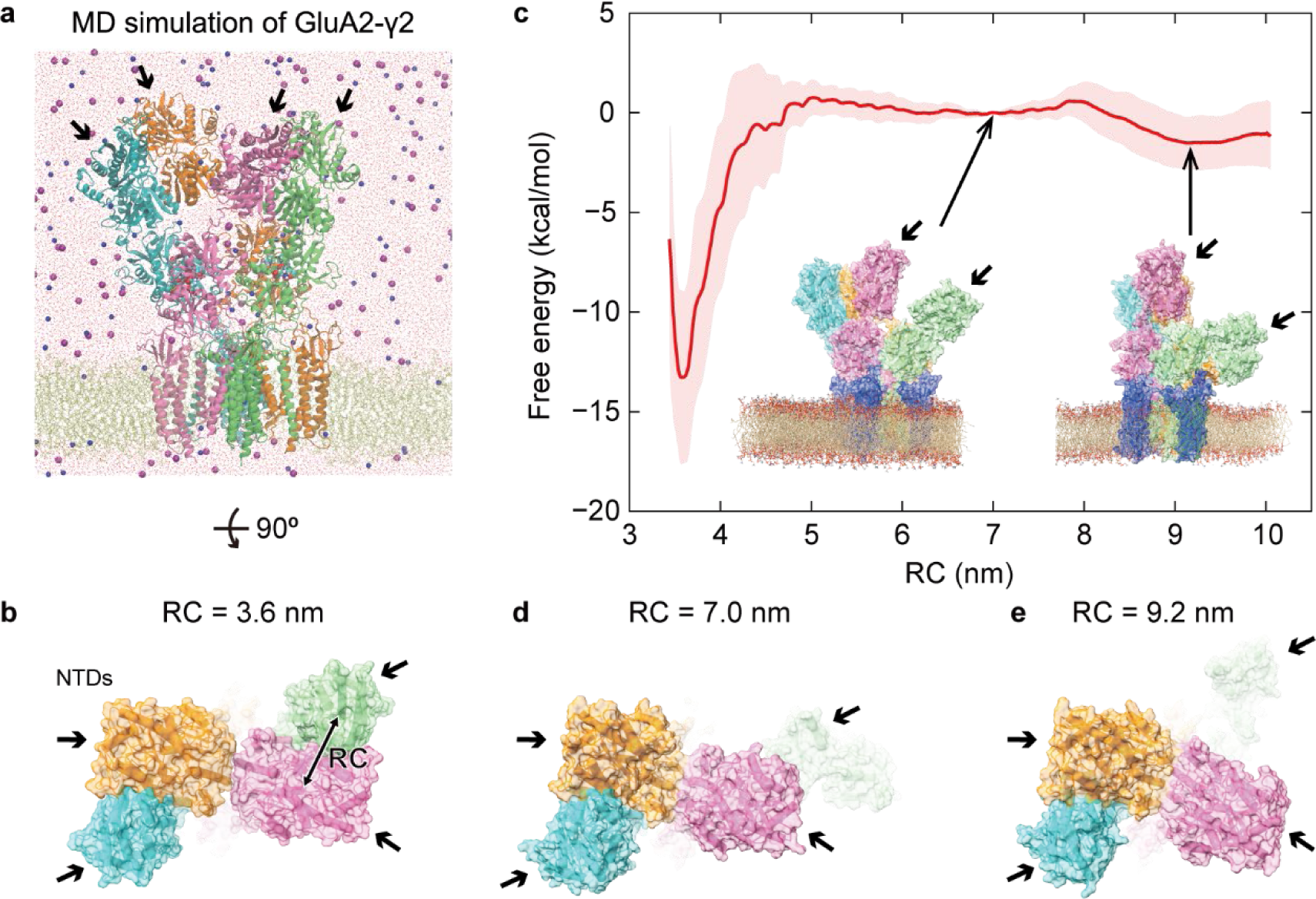
Molecular dynamics simulation of GluA2-γ2 and free energy profile. **a**, The simulation system of GluA2-γ2 with lipids. Blue balls, purple balls, and red dots are Na^+^, Cl^−^, and water molecules, respectively. Glu and CTZ are shown by the spacefill model (see also Supplementary Videos 5 and 6). Black arrows indicate the NTD monomers (**a**-**e**). **b**, Top view of GluA2-γ2. The reaction coordinate (RC) is the centre-to-centre distance between two NTD monomers (green and magenta). **c**, Free energy profile of GluA2-γ2 in lipids as a function of RC. A snapshot of MD simulations at RC = 7.0 nm and 9.2 nm are shown in the inset. The standard deviation (SD) is shown by shading around the mean free energy. **d, e**, Top view of GluA2-γ2 at RC = 7.0 nm (**d**) and 9.2 nm (**e**).

### Intra- and intersubunit exchange between the NTD dimers upon NTD-dimer splitting

Interestingly, we found that after NTD-dimer splitting, the NTDs occasionally reassembled with a different subunit, resulting in intrasubunit dimer exchange (Fig. 6a and Supplementary Video 3). Moreover, when two or more GluA2-γ2 molecules exist in close proximity, they form different combinations of NTD dimers by intersubunit dimer exchange (Fig. 6b and Supplementary Video 7). Previous studies emphasized the importance of the NTD in facilitating the clustering of AMPARs at synaptic sites^36, 37, 52^. Accordingly, we speculated that the intersubunit exchange of the NTD dimers has the potential to contribute to the synaptic clustering of AMPARs.

**Fig. 6:**
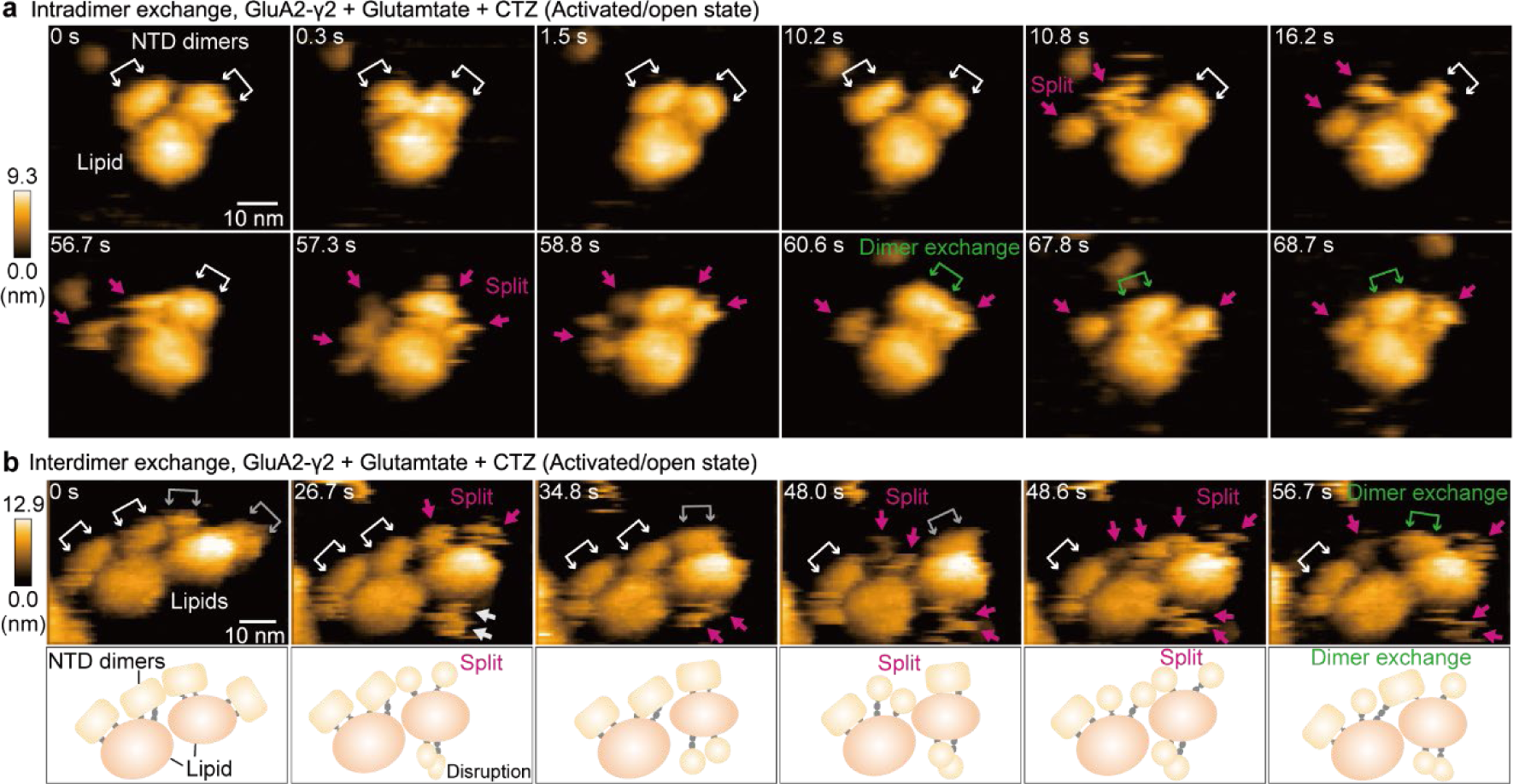
Intra- and interdimer exchange of the NTD dimers in GluA2-γ2. **a, b**, Sequential HS-AFM images of intradimer (**a**) and interdimer (**b**) exchange of GluA2-γ2 in Glu and CTZ (see also Supplementary Video 7). Magenta arrows highlight NTD-dimer splitting. White and grey double-headed arrows indicate the original NTD dimers, while green double-headed arrows indicate the NTD dimers after the interdimer exchange. Schematic drawing of the interpretation of HS-AFM images (**b**, bottom). Imaging parameters: scanning area = 80 × 64 nm^2^ (100 × 80 pixels); frame rate = 3.3 frames/s. The displayed areas are 60 × 50 nm^2^ (**a**) and 65 × 45 nm^2^ (**b**). Z-scale bars indicate height in nm. HS-AFM experiments were repeated at least three times independently with similar results.

### Interactions between NTD dimers and neuronal pentraxin 1

The NTDs are thought to contribute to synaptic clustering by interacting with extracellular scaffold proteins, particularly NP1 (ref.^38, 39^). Therefore, we next investigated the interaction between the NTD and NP1. First, we used AlphaFold2 (ref.^53^) to predict the complex structure of GluA2-γ2 and the pentraxin (PTX) domain of NP1 (NP1_PTX_). The Alphafold2-multimer prediction for the GluA2 NTD dimer and two NP1_PTX_ molecules yielded structural models with high pLDDT and PAE scores (Supplementary Fig. S3a,c). Two NP1_PTX_ molecules bind to the presynaptic side of the GluA2 NTD dimer at the same binding site on each NTD protomer (named site 1 in Fig. 7a). Extensive interactions occur at the interface between them (site 1 in Extended Data Fig. 8a). Interestingly, while the GluA2 residues involved in NP1_PTX_ binding are highly conserved among AMPARs (Extended Data Fig. 8b), some of them are replaced in GluD2, which is the iGluRs family member delta-2 and known to have no interaction with NP1_PTX_ (ref.^54^). For example, Tyr41 of GluA2 and Asp367 of NP1_PTX_ form a hydrogen bond, and Tyr41 is conserved as either tyrosine or histidine among AMPARs (site 1 in Extended Data Fig. 8). In GluD2, the corresponding residue is replaced by aspartate, which prevents the formation of a hydrogen bond with Asp367 of NP1_PTX_. Similarly, Asp63 and Arg83 of GluA2 are replaced by threonine and glutamine in GluD2, respectively (site 1 in Extended Data Fig. 8). These observations reasonably support the plausibility of the predicted structure.

**Fig. 7:**
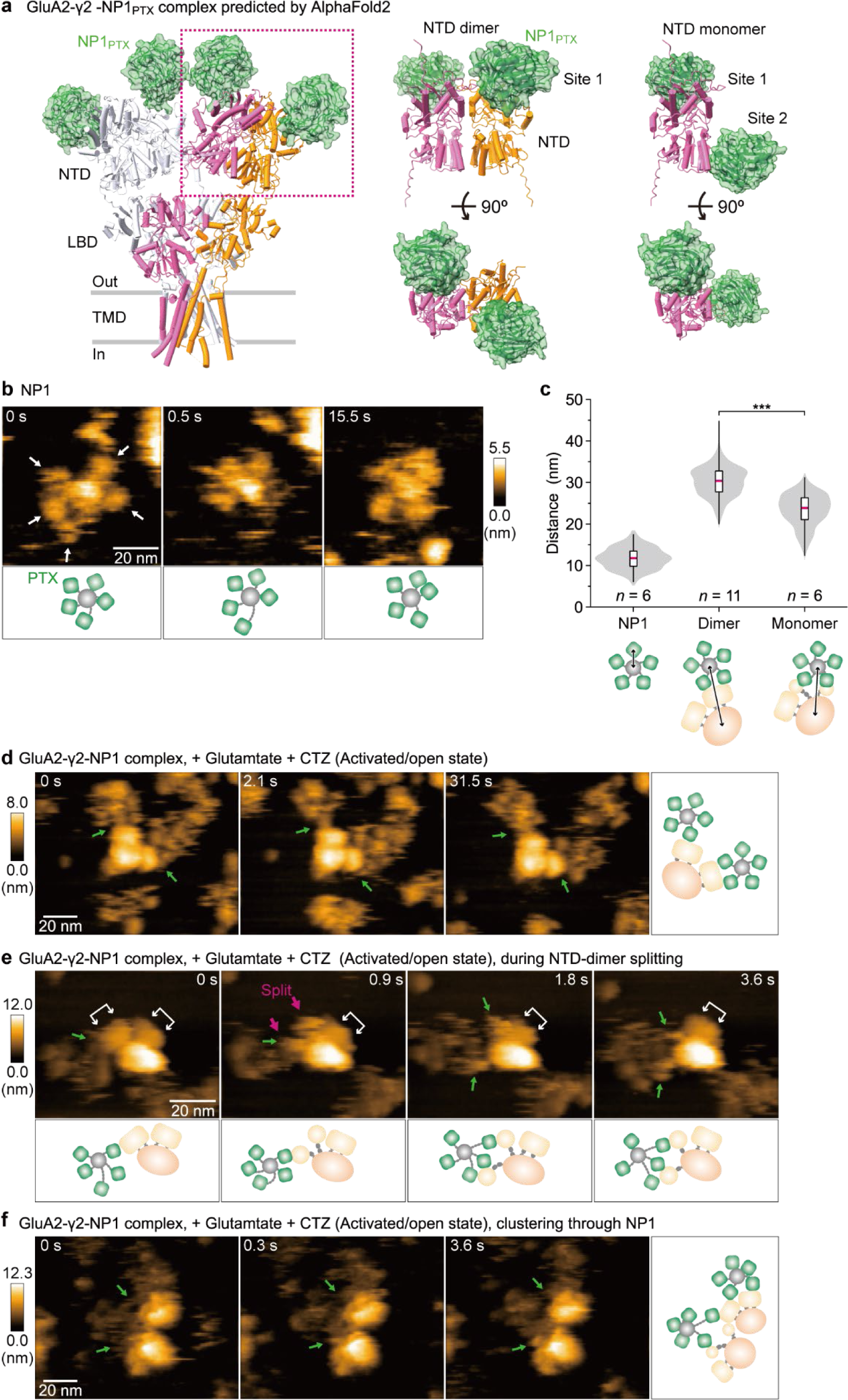
NTD-dimer splitting exposes an additional binding site with neuronal pentraxin 1. **a**, Structures of the GluA2 NTD-NP1_PTX_ complex predicted by AlphaFold2. Right: Predicted binding sites of NP1_PTX_ to the NTD dimer (Site 1) and monomer (Sites 1 and 2). Left: The full-length GluA2 tetramer (PDB: 3KG2) is also shown with its NTDs superimposed on those of the predicted structure. **b, d-f**, Sequential HS-AFM image of NP1 (**b**) and GluA2-γ2 with NP1 in the presence of Glu+CTZ (**d-f**) (see also Supplementary Video 8). White, green, and magenta arrows indicate NP1_PTX_, NP1 bound to the NTDs, and NTD-dimer splitting, respectively. Schematic drawing of the interpretation of HS-AFM images (**b** and **e**, bottom, **d** and **f**, right). Imaging parameters: scanning area = 120 × 96 nm^2^ (240 × 96 pixels) (**b, d, f**) and 80 × 64 nm^2^ (200 × 80 pixels) (**e**); frame rate = 2.0 fps (**b**), 3.3 fps (**d-f**). The displayed areas are 70 × 70 nm^2^ (**b**), 120 × 96 nm^2^ (**d, f**), and 80 × 64 nm^2^ (**e**). Z-scale bars indicate height in nm. HS-AFM experiments were repeated at least three times independently with similar results. **c**, Distances between the centre of NP1 or NP1_PTX_ and the centre of lipids in different NTD oligomeric states. \*\*\**P* < 0.001 (Kruskal‒Wallis test with Mann–Whitney *U* post hoc test).

Furthermore, we directly visualized the GluA2-γ2-NP1 complex using HS-AFM. For this experiment, GluA2-γ2 and NP1 were premixed prior to capturing HS-AFM images. Notably, HS-AFM images of NP1 showed a circular oligomeric structure with a radius of approximately 12 nm, conceivably held together by flexible linkers containing cysteines at the N-terminus region (Fig. 7b,c). Although this configuration of NP1 corroborates the previously expected oligomeric structure^39^, the HS-AFM images further demonstrated that NP1_PTX_ forms a highly mobile structure (Fig. 7b). Additionally, HS-AFM images of the GluA2-γ2-NP1 complex showed that NP1_PTX_ bound to the tips of the NTDs in both the apo and open states (Fig. 7d and Supplementary Video 8).

### NTD-dimer splitting exposed another binding site with NP1

Interestingly, the Alphafold2-multimer prediction for a GluA2 NTD monomer and a single NP1_PTX_ molecule also yielded structural models with high pLDDT and PAE scores (Supplementary Fig. S3b,d) but predicted a distinct binding site from the one we described above (site 1 in Fig. 7a). The new binding site is located at the NTD dimer interface, which is exposed to the solvent side only upon NTD dimer separation (named site 2 in Fig. 7a). Both hydrophilic and hydrophobic interactions are observed at this binding site (Extended Data Fig. 8). As in the case of site 1, while many residues at site 2 are highly conserved among AMPARs (Extended Data Fig. 8b), some of the residues in GluD2 are replaced in a way that disrupts interactions with NP1_PTX_ (Extended Data Fig. 8). For example, Ala177 in GluA2, which contributes to the hydrophobic interactions with Ala368 and Thr369 of NP1_PTX_, is replaced by leucine with a bulkier side chain in GluD2, which is unfavourable for maintaining the interactions. In addition, Glu204 of GluA2 forms a hydrogen bond with the side chain of Tyr238 of NP1_PTX_ but is replaced by a leucine in GluD2 (Extended Data Fig. 8). These consistencies also support the prediction result, as we mentioned above for site 1. Furthermore, 5 μs MD simulations starting from the structures predicted by AlphaFold2 confirmed the stability of these two binding sites, indicating that two NP1_PTX_ molecules can simultaneously and stably bind to the NTD monomer over a microsecond time scale (Extended Data Fig. 9 and Supplementary Video 9).

Subsequently, we used HS-AFM to directly observe the interaction between NP1 and the NTDs during the process of NTD-dimer splitting. Intriguingly, upon NTD-dimer splitting, NP1 protomers within the oligomer, which were initially bound to the NTD dimers, appear to interact with the NTD monomer on the inner surface (presumed to correspond to site 2) (Fig. 7e and Supplementary Video 8). During NTD-dimer splitting, the individual NTD monomers exhibited vigorous and dynamic motion due to their lack of adsorption to the AFM substrate. Consequently, determining the specific binding sites of NP1 on the NTD monomers from the HS-AFM images posed a considerable challenge. Accordingly, considering that site 2 is located on the lipid side of the NTD, we measured the distance between the centre of the NP1 oligomer and the centre of the lipid of nanodiscs containing GluA2-γ2. The results showed that the distance significantly decreased during NTD-dimer splitting (Fig. 7c). This implies that the NP1 oligomer came closer to the lipid region by interacting with an additional binding site on the NTD (site 2). Moreover, we observed two GluA2-γ2 molecules interconnected via an NP1 oligomer (Fig. 7f). This suggests that NP1 oligomers mediate clustering by associating with multiple AMPARs in a bridging configuration. Hence, these findings indicate that NTD-dimer splitting amplifies the number of binding sites between the NTD and the NP1 oligomer, highlighting the potential of AMPARs to promote synaptic clustering.

## DISCUSSION

AMPAR is an ion channel with a crucial function in rapid excitatory neurotransmission in the mammalian central nervous system. Given its great importance, extensive studies have been conducted employing diverse methodologies, such as X-ray crystallography, electron microscopy, and computational simulation, to unravel its molecular mechanism^4^. Despite the abundance of structural information, the structural dynamics of NTDs at the single-molecule level remain elusive. In this study, we employed HS-AFM to observe the conformational dynamics of major functional states of GluA2-γ2 within the lipid environment. Drawing upon our HS-AFM observations, we present the following key findings: (i) elucidation of the structural dynamics of the NTD dimers, (ii) characterization of the increased separation between two NTD dimers in both the open and desensitized states, (iii) identification of the restricted fluctuations of the NTD dimers in the desensitized state (Fig. 3), and (iv) the occurrence of NTD-dimer splitting and subsequent reassembly with different subunits (intersubunit exchange) and increased numbers of binding sites to NP1.

In the activated/open state, the spatial separation between the two NTD dimers increases, and they adopt a conformational transition that takes them further away from the lipid (Fig. 3). In contrast, in the desensitized state, the two NTD dimers remained spatially separated, adopting positions in closer proximity to the lipid, and their fluctuations were restricted (Fig. 3). Recent cryo-EM analyses of the desensitized state of GluA2 in the complex with TARP γ5 and GSG1L have revealed that Quis binding induces a spatial approach of the LBDs towards the lipid. Concurrently, this binding provokes a conformational change of the NTDs to move towards the lipid^47^. These reports are in good agreement with our HS-AFM results. Furthermore, it is conceivable that the fluctuations of the NTD dimers were suppressed by the stronger interaction with the LBD. This conformational clumping of the NTD dimers might inhibit the conformational change of the TMDs, thereby blocking ion permeation. Consistently, the deletion of the NTDs induces decelerated initiation of desensitization and accelerated recovery from desensitization of the agonist responses of AMPARs^30^.

Furthermore, based on our finding of NTD-dimer splitting, we proposed a potential mechanism of the synaptic clustering of AMPARs through NTD dimer exchange and NP1 binding (Fig. 8). AMPAR trafficking to excitatory postsynaptic sites and its synaptic clustering have been investigated for decades because they determine the strength of excitatory synapses^1, 6^. Previous reports have highlighted that in rat hippocampal neurons, AMPARs often localize to synapses in discrete clusters with approximately 70-100 nm in diameter, comprising approximately 20 receptors^55^. These clusters appear to be lined up neatly at presynaptic release sites for fast and efficient synaptic transmission^56, 57^. Assuming these clusters are circular with a radius of 42.5 nm, the derived AMPAR density in the cluster is approximately 0.0035 receptors/nm^2^. In our MD simulations, the density of GluA2-γ2 equated to 0.0033 receptors/nm^2^. A distinct NTD monomer, which emerged after the enforced dissociation of the NTD dimer in the MD simulations, was observed to interact autonomously with the NTD in the neighbouring simulation box, subject to periodic boundary conditions (Extended Data Fig. 10). Although interactions across the simulation boxes are typically considered artefacts, this result underscores that the spatial relationship between the dissociated NTD monomer and the NTD at this density is sufficiently close to form the NTD dimer through interdimer subunit exchange. Thus, MD simulations also supported that the NTD dimer subunits can undergo exchange at densities consistent with those of AMPAR clusters in neurons. In addition, recent cryo-EM analyses of GluA1/TARPγ3 and GluA2(F231A)/TARPγ2 revealed that approximately 20% of particles deviate from the canonical AMPAR architecture, notably missing the domain swapping between the NTD and LBD dimers^49^. If the transition between swapping and nonswapping of the NTD dimers occurs, this would support our HS-AFM finding that demonstrated NTD-dimer splitting.

**Fig. 8:**
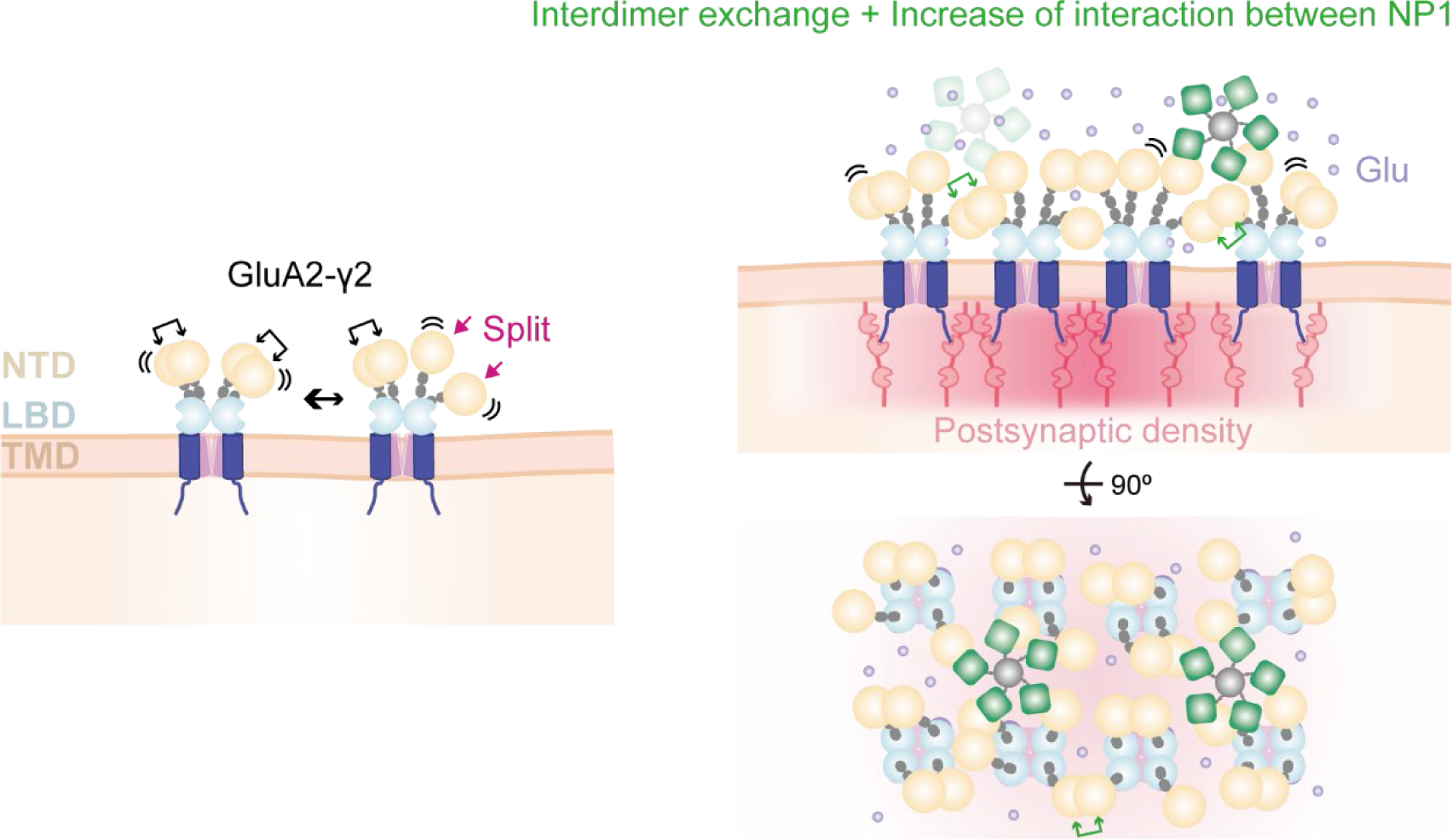
Proposed model outlining the role of the NTD in the synaptic clustering of AMPARs. The NTD dimers of GluA2-γ2 fluctuate thermally, changing the distance between them and sometimes splitting. Upon reaching the postsynaptic density, GluA2-γ2 is mainly localized by the interaction of PSD-95 with the C-terminus of TARPs. Simultaneously, when NTD-dimer splitting occurs, the NTDs interact with neighbouring GluA2-γ2 molecules by interdimer exchange and NP1 binding to form clusters (right). Since Glu-bound GluA2-γ2 in the open state has a longer lifetime in the splitting state of the NTD dimers, AMPAR clustering may be enhanced at synaptic active sites where Glu is released.

The interaction between the NTD and presynaptic proteins (neuronal pentraxins) is important for the synaptic clustering of AMPARs^35, 38–40^. Our structural predictions showed that the number of binding sites to NP1 increases upon the initiation of NTD-dimer splitting. In the NTD dimer, NP1 associates with a singular binding site, which is distal from the lipid. However, in the NTD monomer, an additional binding site for NP1 is unveiled at the dimer-forming interface, which increases the affinity between the NTD and NP1. Notably, in the activated/open state in the presence of Glu+CTZ, the lifetime of the NTD splitting state was longer than that of the apo state (Fig. 4). Consequently, in synaptic regions where glutamate is released, the NTDs tend to transition towards the monomer state, which further strengthens the binding to NP1 (Fig. 8).

Notably, the optimization of the experimental conditions of HS-AFM, in particular the incorporation of a fluorochemical surfactant in the imaging buffer, enables lateral observation of the extracellular domain of membrane proteins. This method can potentially be applied to other membrane proteins to visualize the dynamics of large domains protruding from lipids, which are frequently encountered in eukaryotic systems. Such observations can greatly contribute to unravelling the functional implications associated with their dynamics.

Other isoforms of AMPARs would be suitable examples for the application of HS-AFM presented here. AMPAR tetramers have multiple isoforms^58, 59^ and are heterotetramers of GluA1-GluA4 subunits in vivo^60–62^. Given the importance of GluA2 as a pivotal subunit that predominantly occurs in native AMPAR tetramers, our present study used HS-AFM to examine AMPARs containing a single GluA2 with TARP γ2. In future research, further observations of heterotetramers involving various subunits have the potential to deepen our understanding of the intricate mechanism of clustering under near-physiological conditions. Furthermore, the application of our method to other glutamate receptors, including NMDA receptors, kainate receptors, and metabotropic glutamate receptors, is also important for understanding the interplay between functions and structural dynamics.

### Conclusion

HS-AFM provided insights into the structural dynamics of the NTD dimers of GluA2-γ2 within the lipid environment. Specifically, the NTD dimers fluctuated horizontally relative to the lipid edge. The range of fluctuations between dimeric NTDs increased in the open state, while their frequency was restricted in the desensitized state. Although further experimental evidence is necessary to confirm the occurrence of NTD-dimer splitting at the cellular level, our HS-AFM observations of the interdimer exchange of the NTD, coupled with augmented affinity to NP1, suggest a new hypothesis regarding glutamate-stimulated AMPAR clustering at synaptic sites, which enables rapid neurotransmitter function mediated by AMPARs.

## Methods

### Expression and purification of GluA2-γ2

HEK293S GnTIˉ cells (ATCC CRL-3022) were cultured in FreeStyle™ 293 Expression Medium (Gibco, 12338018) containing 1% fetal bovine serum (FBS), incubated in 5% CO_2_ at 37°C, and routinely passaged every three days.

The expression construct of *Rattus norvegicus* GluA2 fused to TARP γ2, originally used for single particle cryo-EM analysis of GluA2-γ2 (ref.^23^), was synthesized by Genewiz (Suzhou, China) and was subcloned and inserted into the pEG BacMam vector plasmid^63^ with an octahistidine tag, EGFP, and a tobacco etch virus (TEV) protease cleavage site at its C-terminus. In the GluA2-γ2 fusion construct, the C-terminus of GluA2 was fused to the N-terminus of TARP γ2 via a glycine-threonine linker^23^.

DH10Bac *E. coli* cells (Gibco, 10361012) were transformed with the plasmid for bacmid preparation, and Sf9 cells were transfected with the bacmid using FuGENE HD (Promega, E2311). Then, recombinant baculovirus was used to infect HEK293S GnTIˉ cells. Next, 10 mM sodium butyrate was added to induce GluA2-γ2 expression. Cells were harvested by centrifugation (5,400 × g, 10 min), disrupted by sonication, and then further centrifuged (7,600 × g, 20 min). The supernatants were solubilized with solubilization buffer (20 mM Tris-HCl pH 8.0, 150 mM NaCl, 2% n-dodecyl-beta-D-maltopyranoside (DDM), 0.4% cholesteryl hemisuccinate (CHS), 30 mM imidazole, 1 mM phenylmethylsulfonyl fluoride (PMSF), 5.2 μg/mL aprotinin, 2 μg/mL leupeptin, and 1.4 μg/mL pepstatin A) for 2 hours at 4°C and then ultracentrifuged to remove insolubilized materials (200,000 × g, 1 hour). Supernatants were applied to an open column loaded with TALON beads (Cytiva, 28957502). The beads were washed with Wash Buffer 1 (20 mM Tris-HCl pH 8.0, 150 mM NaCl, 0.05% DDM, 0.01% CHS, 30 mM imidazole, 10 mM MgCl_2_, 1 mM ATP) to remove the chaperone and then with Wash Buffer 2 (20 mM Tris-HCl pH 8.0, 150 mM NaCl, 0.05% DDM, 0.01% CHS, 30 mM imidazole). GluA2-γ2 proteins were eluted with elution buffer (20 mM Tris-HCl pH 8.0, 150 mM NaCl, 0.05% DDM, 0.01% CHS, 250 mM imidazole) and then dialyzed against dialysis buffer (20 mM Tris-HCl pH 7.5, 150 mM NaCl, 0.05% DDM, 0.01% CHS) after the addition of TEV protease to digest the His-EGFP tag. GluA2-γ2 proteins were loaded onto a Superose 6 Increase 10/300 GL column (Cytiva, 29091596) equilibrated with gel filtration buffer (20 mM HEPES pH 7.0, 150 mM NaCl, 0.05% DDM, 0.01% CHS). The major peak fractions were collected, concentrated to 1 mg/mL using an Amicon Ultra Concentrator (Millipore, UFC810024) and stored at −80°C (Extended Data Fig. 2a). To verify the monodispersity of purified GluA2-γ2, 1 µg of purified GluA2-γ2 was applied to a Superdex 200 Increase 10/300 GL column (Cytiva, 228990944) equilibrated with FSEC buffer (50 mM Tris-HCl, pH 8.0, 150 mM NaCl, 0.03% DDM) for Trp-based FSEC (excitation: 280 nm, emission: 325 nm) (Extended Data Fig. 2b)^64, 65^.

The expression construct of GluA2 (ref.^11^) not fused to TARP γ2 was synthesized by Genewiz (Suzhou, China) and then subcloned and inserted into the pFastBac vector plasmid with an octahistidine tag, EGFP, and a human rhinovirus 3C protease (HRV3c) cleavage site at its C-terminus. Sf9 cells were infected with recombinant baculovirus to express GluA2. The GluA2 protein was purified by a similar procedure to the purification of GluA2-γ2 described above, except that CHS was not used for purification and HRV3c protease was used instead of TEV protease to remove the His-EGFP tag.

### Reconstitution of GluA2-γ2 into a lipid

GluA2-γ2 reconstitution into a lipid bilayer was achieved using the nanodisc method with slight modifications^44^ based on the manufacturer’s protocol (Sigma‒Aldrich, USA). Initially, soy phospholipids (120 μg) obtained from soybean (#11145, Sigma‒ Aldrich) were dissolved in chloroform and evaporated under N_2_ gas to completely remove the solvent. The resulting soy phospholipid was then suspended in 50 μL of buffer containing 20 mM HEPES-NaOH (pH 7.4), 100 mM NaCl, and 4% DDM. Sonication with a tip sonicator (TAITEC, Japan) was performed for approximately 1 min. To this soy phospholipid suspension, solubilized GluA2 or GluA2-γ2 (1 nmol) and membrane scaffold proteins (50 μL, final concentration 1 mg/mL) (MSP1E3D1, Sigma‒ Aldrich) were added, and the mixture was rotated at 4°C for approximately 1 hour. Subsequently, 60 mg of Bio-Beads SM-2 (#1523920, Bio-Rad) was added, and the samples were dialyzed overnight at 4°C in the presence of detergent. Although the manufacturer’s protocol recommends further fractionation and size purification of the protein-embedded nanodisc sample to achieve a diameter of approximately 10 nm, in our HS-AFM experiments, we did not perform this purification step. Instead, we obtained flat membranes less than 30 nm in diameter.

### Preparation of neuronal pentraxin 1 protein

Recombinant Human Neuronal Pentraxin 1 Protein (NP1) was purchased from R&D systems (#: 7707-NP, bio-techne, USA).

### HS-AFM apparatus

HS-AFM observations were performed using a custom-designed HS-AFM instrument^42, 44, 66^. In tapping mode, an optical beam deflection detector was employed, utilizing a 780 nm, 0.8 mW infrared (IR) laser to detect the deflection of the cantilever (BL-AC10DS-A2, Olympus, Japan). The IR beam was focused onto the back side of the cantilever through a 60× objective lens (CFI S Plan Fluor ELWD 60X, Nikon, Japan). The reflection of the IR beam from the cantilever was captured by a two-segmented PIN photodiode (MPR-1, Graviton, Japan). The spring constant of the cantilever was approximately 100 pN/nm. In a liquid environment, the cantilever exhibited a resonant frequency of approximately 400 kHz and a quality factor of approximately 2. The original AFM tip had a triangular section resembling a bird beak. To further improve spatial resolution, an amorphous carbon tip was grown on the bird beak tip by electron beam deposition (EBD) using a scanning electron microscope (FE-SEM; SUPRA40VP, Carl Zeiss, Germany). The additional AFM tip had a length of approximately 500 nm, and the tip apex had a radius of approximately 1 nm after further plasma etching with a plasma cleaner (Tergeo, PIC Scientific, USA). All HS-AFM images were acquired using the cantilever with the additional AFM tip. The free oscillation amplitude of the cantilever was less than 1 nm, and the set-point amplitude during HS-AFM observations was adjusted to approximately 90% of the free amplitude. To minimize the interaction force between the sample and the AFM tip, HS-AFM was operated in “only trace imaging” (OTI) mode^67^. HS-AFM operation and data collection were performed using laboratory-developed software based on Visual Basic. NET (Microsoft).

### HS-AFM observations

HS-AFM observations of GluA2 and GluA2-γ2 in the apo state were performed on mica in buffer A (50 mM Tris-HCl, pH 8.0, 150 mM NaCl, 10% glycerol), with or without 0.1% (1H, 1H, 2H, 2H-perfluorooctyl)phosphocholine (O310F, Anatrace, USA). HS-AFM observations of GluA2-γ2 in the closed state were performed on mica in buffer A supplemented with 0.1% O310F and 0.3 mM ZK200775 (#2345, Tocris, UK). Similarly, HS-AFM observations of GluA2-γ2 in the open state were performed on mica in buffer A with 0.1% O310F, 3 mM L-glutamate (Glu, #0218, Tocris, UK) and 0.1 mM cyclothiazide (CTZ, #0713, Tocris, UK). Similarly, HS-AFM observations of GluA2-γ2 in the desensitized state were performed on mica in buffer A with 0.1% O310F and 1 mM L-quisqualate (Quis, #0188, Tocris, UK). For HS-AFM observations of the GluA2-γ2 and NP1 complex, GluA2-γ2 and 25 nM NP1 were premixed in buffer A with or without 3 mM Glu and 0.1 mM CTZ and incubated at room temperature for 5 min. Then, HS-AFM observations were performed in buffer A with 0.1% O310F and with or without 3 mM Glu and 0.1 mM CTZ. All HS-AFM experiments were performed at room temperature (24– 26°C) and repeated independently at least three times, yielding consistent results.

### HS-AFM image processing and data analysis

HS-AFM images were processed using Fiji (ImageJ) software (NIH, USA)^68^. To reduce noise, a mean filter with a radius of 0.5 pixels was applied to each HS-AFM image. The Template Matching and Slice Alignment plugin for ImageJ was used to correct for drift between sequential images. To track the coordinates of the centres of the NTD, LBD, lipid of GluA2-γ2, NP1_PTX_, and NP1 in all HS-AFM experimental conditions, either the TrackMate^69^ or MtrackJ^70^ plugins for ImageJ were utilized semimanually.

The analysis involved measuring the distance between the NTD dimers of GluA2-γ2 in apo (3887 frames of 16 molecules), ZK200775-bound (1676 frames of 7 molecules), Glu- and CTZ-bound (3713 frames of 14 molecules), and Quis-bound (2709 frames of 9 molecules) forms (Fig. 2c). Additionally, the distance between the NTD dimers and the lipid edge in various experimental conditions was as follows: apo (220 frames of 11 molecules), ZK200775-bound (142 frames of 7 molecules), Glu- and CTZ-bound (202 frames of 10 molecules), and Quis-bound (202 frames of 10 molecules) forms (Fig. 2d). To compare distances across different HS-AFM experimental conditions, the distances were normalized by considering the median length of the NTD dimer in 10 arbitrary frames from each sequential HS-AFM image.

To determine the sum of squared deviations, we computed the deviation between the distances of the NTD dimers in each frame, squared those values, and then summed them for each frame (*n* = 12 (apo), 7 (+ZK), 11 (+Glu+CTZ), and 6 (+Quis) molecules). Subsequently, we compared the sum of squared deviations at 60 s for each experimental condition and obtained statistical differences (Dunnett’s test, Extended Data Fig. 5).

The distances between the centre of NP1 and NP1_PTX_ (156 frames of 6 oligomers), the lipid centre of the nanodisc with embedded GluA2-γ2 in the NTD dimer (1562 frames of 11 molecules) and the NTD monomer (694 frames of 6 molecules) were analysed (Fig. 7c).

### Analysis of trajectories

To quantify the moving range of the NTDs, the LBDs and the lipids in GluA2, we computed the gyration radius, *R*_*g*_, as the root-mean-square displacement of their positions from the average position^71^.

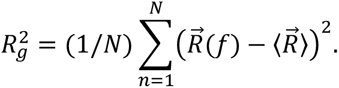

where 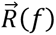 represents the positions of the trajectories of the NTDs, the LBDs, and the lipids in frame f. 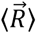 denotes the average position of the corresponding trajectories. *N* corresponds to the number of frames. For the analysis of *R_g_*, GluA2 (*n* = 6) was analysed (Extended Data Fig. 1).

### AFM simulation

BioAFMviewer software^72^ was used to validate the observed topographies of GluA2-γ2 by HS-AFM. The simulation involved nonelastic collisions of a rigid cone-shaped tip model with the rigid van der Waals atomic model of the protein structure^73^. For the structural template, two subunits of GluA2-γ2 were extracted from PDB:5KBU. The simulation parameters were as follows: scan step (0.81 nm), tip probe sphere radius (3.6 nm), and cone half-angle (20°) (Extended Data Fig. 3).

### Correlation function of NTD-dimer splitting

The correlation function (*C*(*t*)) is defined by the following equation^74^:

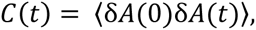

where the angular bracket represents the ensemble and time averaging. Here, δ*A*(*t*) = *A*(*t*) − 〈*A*〉, with *A*(*t*) equal to 1 when the NTD dimers are split and 0 otherwise. *C*(*t*) was fitted using triple exponential functions: A_1_exp(−*t*/τ_1_) + A_2_exp(−*t*/τ_2_) + A_3_exp(−*t*/τ_3_), where A_1_ + A_2_ + A_3_ = 1. The lifetime from stable to split states (τ_01_) was calculated using the equation:

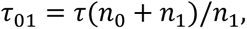

where t represents t_1_, t_2_, or τ_3_, and *n*_1_ and *n*_0_ are the numbers of frames in which the dimers are split and not split, respectively. Similarly, the lifetime from split to stable states (T_10_) was calculated as

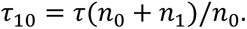

The correlation function of NTD-dimer splitting in GluA2-γ2 was analysed in apo (6042 frames of 18 molecules (36 dimers)), ZK200775-bound (3065 frames of 8 molecules (16 dimers)), Glu- and CTZ-bound (6418 frames of 20 molecules (40 dimers)) and Quis-bound (3544 frames of 11 molecules (22 dimers)) forms.

### AlphaFold2 prediction

The predicted structural models of the GluA2 NTD (residues 21-404, P19491) in complex with the neuronal pentraxin 1 (NP1) pentraxin (PTX) domain (residues 225-429, Q15818) were generated using AlphaFold2 (v2.3.1)^53^ and ColabFold (v1.5.3)^75^ (Supplementary Fig. S3). The predicted structural models with the highest scores were selected for further analysis (Supplementary Data 1 and 2). PyMOL software (https://pymol.org/) was used to generate all structure figures. Clustal Omega^76^ and ESPript 3.0 (ref.^77^) were used to generate the sequence alignment figure (Extended Data Fig. 8).

### Molecular dynamics (MD) simulation of GluA2-γ2 and calculation of the free energy profile

GluA2-γ2 with Glu and CTZ (PDB code: 5WEO) was embedded in the preequilibrated POPC membrane with approximately 150 mM NaCl solution. Before embedding, the bad contacts in the crystallographic structure were removed by 1,000-step minimization. Amber ff19SB (ref.^78^), Lipid17 (ref.^79^), TIP3P (ref.^80^), the Horn model^81^, and the Joung-Cheatham^82^ models were employed for GluA2-γ2, the POPC lipid molecule, water, Glu, and ions, respectively. The force field of CTZ was developed by a conventional method: the charges on CTZ were calculated by the *ab initio* calculation using Gaussian09 (ref.^83^) at the HF level with a 6-31G* basis set followed by RESP assignment using the antechamber module in AMBER22 (ref.^84^). Atom types of CTZ were assigned by the antechamber module, and the general AMBER force field^85^ was used for CTZ. All the charges and atom types of CTZ are included in the electronic supplement file of the AMBER library file (CTZ.lib in Supplementary Data 1).

The system comprised one GluA2-γ2, 887 POPC molecules, 161,547 water molecules, four Glu, four CTZ, 438 Na^+^ ions, and 438 Cl^−^ ions. The histidine residues were assumed to be in the protonated state at pH = 7, and two cysteine residues were assumed to have a disulfide bond when they were close together. The total number of atoms in the system was 666,719. This information with a box size of periodic boundary conditions is tabulated (Supplementary Table S2).

The MD simulation was performed for 1 ns with constant temperature (300 K) and pressure conditions (1 bar), where harmonic restraints were imposed on all atoms in GluA2-γ2, Glu, and CTZ with a force constant of 2 kcal/mol/Å^2^. In the following 2 ns simulation, no restraints were imposed. In this 3 ns simulation, the gap between the GluA2-γ2 and POPC membranes made in the embedding process was filled, and the membrane area was equilibrated. The phosphate atoms of one-half of lipid molecules in the lower leaflet membrane (opposite to the NTDs and the LBDs) were weakly constrained (with a force constant of 0.02 kcal/mol/Å^2^) to their centre throughout all simulations to suppress unwanted undulation^86^. All MD simulations were performed using AMBER22 (ref.^27^). A Berendsen thermostat^87^ was used to control the temperature, and a Monte Carlo barostat with anisotropic scaling was used to control the pressure. The SHAKE algorithm^88^ was used to keep the bond length having H atoms constant, enabling the use of a time step of 2 fs. The periodic boundary condition was imposed, and long-range interactions were calculated by the particle mesh Ewald method^89^ with a 10 Å cut-off in real space.

Then, to fully equilibrate the system, the MD simulation was performed for 500 ns with constant temperature at 300 K and volume conditions using the Berendsen thermostat^87^. After 200 ns, the potential energy of the system fluctuated, but its average value did not change significantly. Glu and CTZ remained at the binding positions observed in the crystallographic structure. Although the overall structure of GluA2-γ2 was maintained, the NTDs and the LBDs fluctuated slightly in the bulk solution (Supplementary Video 5). The root mean square displacement (RMSD) from the crystallographic structure of alpha carbon atoms in the NTD was less than 0.25 nm and did not increase with time, and the root mean square fluctuations (RMSF) of alpha carbon atoms in the NTD were mostly less than 0.1 nm (Supplementary Fig. S1).

To explore the possibility of NTD-dimer splitting, force was applied between the NTD monomers to split the NTD dimer. The reaction coordinate (RC) was defined by the distance from a centre of the NTD monomer to a centre of the other NTD monomer, where the centre is the averaged position of alpha carbon atoms every 5 residues in the NTD starting from Asn25. RC is approximately 3.6 nm in the crystallographic structure. A harmonic potential was applied with a force constant of 1 kcal/mol/Å^2^ and an equilibrium distance of 10.0 nm. The applied force was proportional to the difference from the equilibrium distance and approximately 4.4 nN when RC = 3.6 nm. NTD-dimer splitting occurred in 10 ns, and the simulation was performed for 500 ns to stabilize the split structure. The RMSF of alpha carbon atoms in the NTD after splitting was very similar to that before splitting (Supplementary Fig. S2), meaning that the NTD dimer split into monomers without breaking. Then, the harmonic potential to make the split structure was turned off, and a 200 ns simulation was performed. The split NTD monomers did not return to the dimer, and one monomer spontaneously bound to the NTD across the periodic boundary condition despite the absence of an attractive external force applied between the monomer and the NTD (A and B in Extended Data Fig. 10). Note that when the equilibrium distance was 5.5 nm, the split monomers immediately (in less than 10 ns) resumed the dimer conformation.

The free energy profile along the RC was obtained using umbrella sampling^90^. The RC was varied from 3.4 nm to 10.0 nm with a window spacing of 2 Å, resulting in a total of 34 windows. A 10 ns equilibrium simulation was performed, followed by a 100 ns sampling simulation. The RC was recorded every 100 fs, so 1 million RC values were sampled at a certain RC. After equilibration, the centre of the harmonic potential was successively shifted from 3.4 nm to 10.0 nm and from 10.0 nm to 3.4 nm. The force constant was 4 kcal/mol/Å^2^ for 3.4 nm ≤ RC ≤ 5.8 nm, where there is a deep free energy well, and 1 kcal/mol/Å^2^ for RC ≥ 6.0 nm. For 3.4 nm ≤ RC ≤ 5.8 nm, the calculations were repeated using the force constant of 2 kcal/mol/Å^2^. The averaged free energy profile was estimated by repeating these calculations three times. The standard deviation of the free energy profile was calculated by these six runs for 3.4 nm ≤ RC ≤ 5.8 nm and three runs for RC ≥ 6.0 nm. Accordingly, the total simulation time was 31.02 μs, and 282 million RC values were sampled in total. The weighted histogram analysis method^91^ was used to construct the free energy profile.

### MD simulation of NTD and NP1_PTX_

To explore the potential binding between NTD and NP1_PTX_, MD simulations were performed for the NTD-NP1_PTX_ systems. The initial structures of the NTD monomer complexed with one NP1_PTX_ molecule and the NTD dimer complexed with two NP1_PTX_ molecules were constructed using AlphaFold2 (ref.^53^) (Fig. 7, Extended Data Fig. 8 and Supplementary Data 2, 3). The initial structure of the NTD monomer with two NP1_PTX_ molecules at sites 1 and 2 was generated by combining the above two structures. Specifically, the NTD dimer with two NP1_PTX_ molecules (both bound at site 1) was taken as the NTD monomer with one NP1_PTX_ molecule (chain B [NTD] and C [NP1_PTX_] in Supplementary Data 3). Then, residues 130 to 190 of the NTD monomer complexed with NP1_PTX_ at site 2 (Supplementary Data 2) were replaced with the corresponding residues from the taken NTD monomer complexed with one NP1_PTX_ at site 1, including one NP1_PTX_ bound to site 2.

The structures generated by AlphaFold2 were optimized by 10,000-step minimization and then dissolved in 150 mM NaCl solution. The system sizes were equilibrated by 1 ns constant temperature (300 K) and pressure (1 bar) simulation. Three replicas were made for three systems, and 5 μs simulations were performed at constant temperature (300 K) and volume conditions. The total simulation time was 45 μs. The simulation conditions, such as the force field and the temperature control method, were the same as those described above. Information on the systems with a box size of the periodic boundary conditions is tabulated in Supplementary Table S3. All data on distances between NTD and NP1_PTX_ were provided in Supplementary Data 4.

### Quantification and statistical analysis

Statistical analyses were performed using Igor Pro9 software (WaveMetrics, USA) and OriginPro 2023 software (OriginLab, USA). A significance level of α = 0.05 was applied for all analyses, and *P* values were adjusted for multiple comparisons when necessary. Unless otherwise stated, the normality of distributions was assessed using the Shapiro‒ Wilk test, and the equality of variance was examined using Bartlett’s test. If the distribution was acceptable, one-way ANOVA was performed, and if it failed, the Kruskal‒ Wallis test was used for analysis of multiple groups with a single independent variable. The Tukey HSD, Dunn, Mann–Whitney *U,* and Dunnett tests were used as follow-up tests for one-way ANOVA or the Kruskal‒Wallis test when comparing every mean to a control mean. A *P* value > 0.05 was not considered statistically significant (N.S.), and *P* values < 0.05 were considered to indicate statistical significance. *P* values of *P* < 0.05 (*), *P* < 0.01 (**), and *P* < 0.001 (^∗∗∗^) are indicated accordingly. The specific tests used for each dataset are described in the figure legends. Data are represented as the mean values ± SDs.

## SUPPLEMENTAL INFORMATION

Supplemental information can be found online at XXXX.

## ACKNOWLEDGEMENTS

This research was supported by the World Premier International Research Center Initiative (WPI), MEXT, Japan (to M.S.), JSPS KAKENHI grant numbers JP20K21122 (to M.S.) and JP21H01771 (to M.S.), the Mochida Memorial Foundation for Medical and Pharmaceutical Research (to M.S.), the Uehara Memorial Foundation (to M.S.), the National Natural Science Foundation of China (32071234, 32271244 and 32250610205 to M.H.), an innovative research team of high-level local universities in Shanghai and a key laboratory program of the Education Commission of Shanghai Municipality (ZDSYS14005 to M.H.), and JST/CREST (JPMJCR1762 to N.K. and H.F.). We thank Prof. Toshio Ando at Kanazawa University for providing the HS-AFM-related apparatus and Youko Mikami, Kaito Onodera, and Yui Okada at Kanazawa University for collecting and analysing the HS-AFM data. Calculations were partly conducted on a supercomputer at the Research Center for Computational Science in Okazaki, Japan (Project: 23-IMS-C101 to T.S.).

## AUTHOR CONTRIBUTIONS

Conceptualization, M.H. and M.S.; methodology, K.U., N.K., and M.S.; software, T.S., H.F., and M.S.; validation, A.S., T.S., and M.S.; formal analysis, A.S., T.S., and M.S.; investigation, A.S., T.S., Y.Z., M.H., and M.S.; resources, Y.Z., T.S., M.H., and M.S.; data curation, M.H., and M.S.; writing – original draft, M.S.; writing – review & editing, all authors; visualization, T.S., M.H., and M.S.; supervision, H.M., and M.S.; project administration, M.S.; funding acquisition, M.H., and M.S.

---

**Extended Data Fig. 1:**
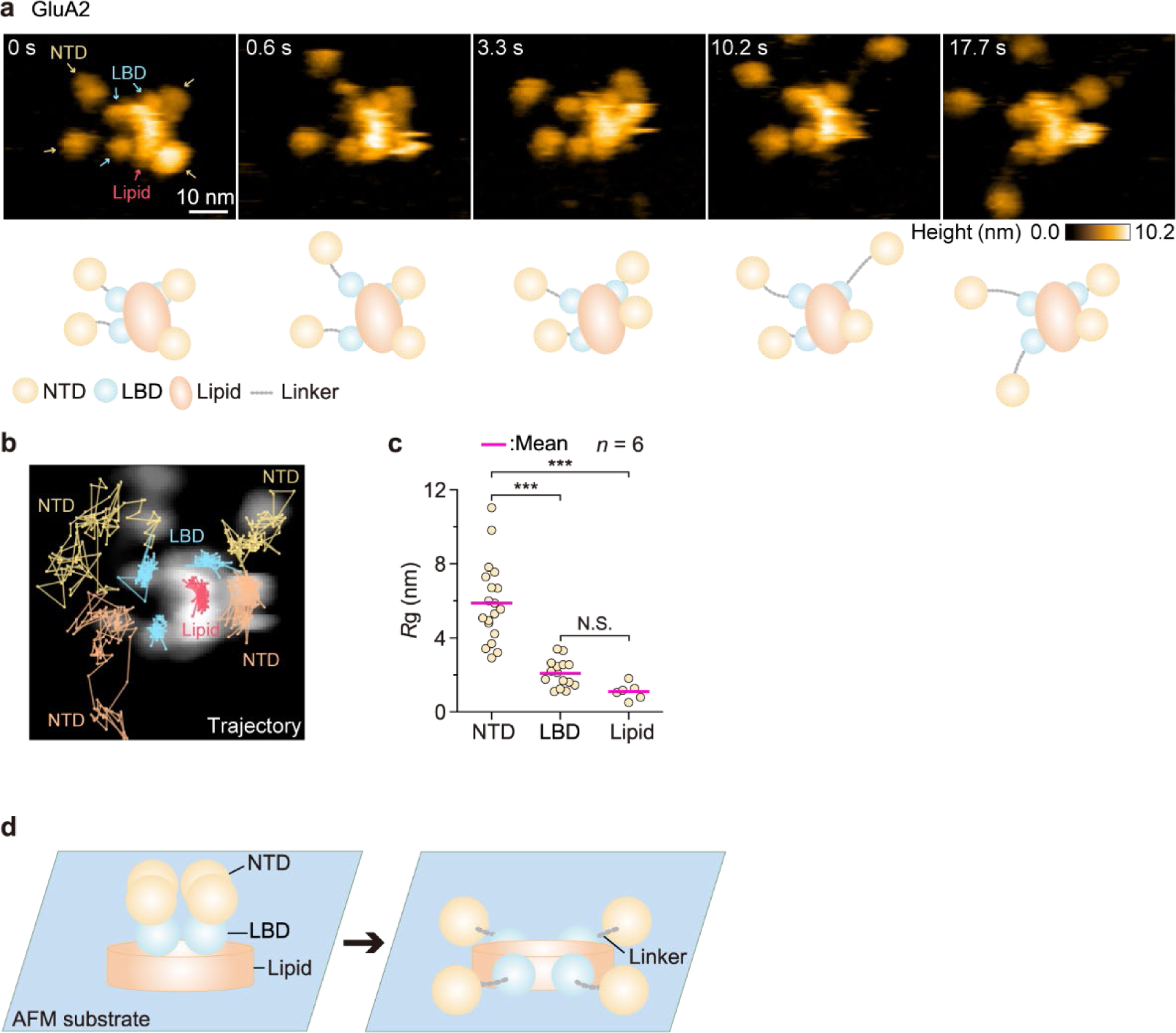
HS-AFM observations of GluA2 alone without O310F on mica, related to Fig. 1. **a**, Sequential HS-AFM images of GluA2 without O310F. Yellow, blue and red arrows indicate the NTDs, LBDs and a lipid, respectively. Diagrams of the presumed structures of the linker in GluA2 are also shown at the bottom of each HS-AFM image. Imaging parameters: scanning area = 60 × 48 nm^2^ (120 × 96 pixels); frame rate = 3.3 frames/s. The displayed area is 60 × 48 nm^2^. HS-AFM experiments were repeated at least three times independently with similar results. **b**, Trajectories of the NTDs, LBDs, and a lipid were tracked for ∼40 s. **c**, *R*_g_ for the NTDs, LBDs, and lipids of GluA2 without O310F (*n* = 6 molecules). not significant (N.S.); \*\*\**P* < 0.001 (one-way ANOVA with Tukey’s HSD post hoc test). **d**, Schematic diagram of the adsorption of GluA2-embedded nanodiscs on a mica surface without O310F. Under these experimental conditions, a predicted GluA2 conformation (left) collapsed on the mica surface; the four NTDs and LBDs moved far apart and independently (right).

**Extended Data Fig. 2:**
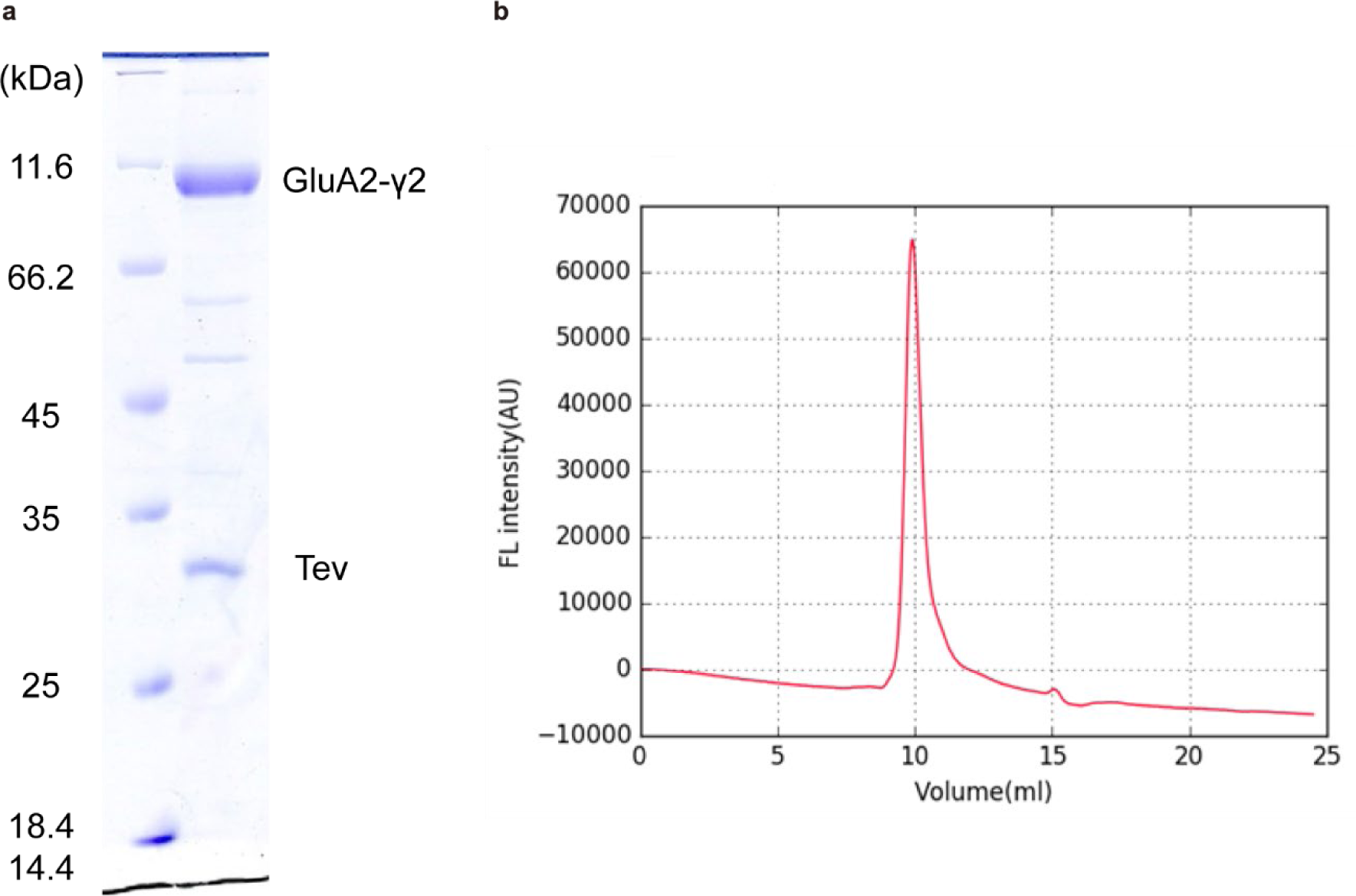
Purified GluA2-γ2 sample, related to Fig. 1. **a**, SDS‒PAGE of purified GluA2-γ2. **b**, FSEC trace of purified GluA2-γ2, as detected by Trp fluorescence.

**Extended Data Fig. 3:**
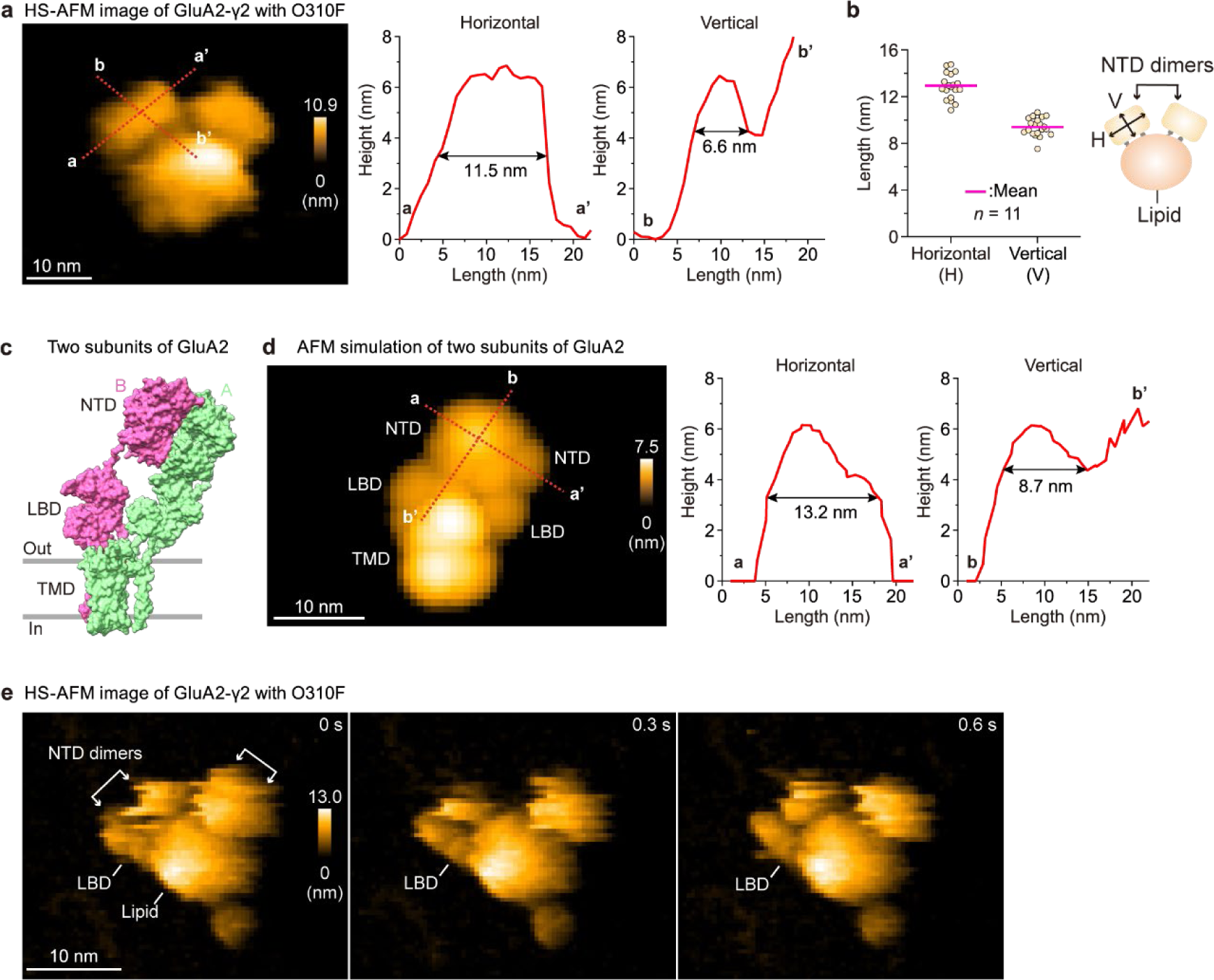
Comparison of HS-AFM and AFM simulation images of GluA2-γ2, related to Fig. 1. **a**, Representative HS-AFM image of GluA2-γ2 with O310F. The cross sections at the red dotted lines a-a’ (horizontal) and b-b’ (vertical) are shown on the right (also in **d**). **b**, Horizontal and vertical lengths of the NTD dimers of GluA2-γ2 with O310F obtained from HS-AFM images (*n* = 11 molecules). **c**, The cryo-EM structure of GluA2-2xγ2 (PDB:5KBU). For comparison with HS-AFM images, only two subunits within tetrameric GluA2 are shown. **d**, A simulated AFM image of two subunits of GluA2-γ2 from **c**. **e**, Sequential HS-AFM images of GluA2-γ2 with O310F. Imaging parameters: scanning area = 80 × 64 nm^2^ (100 × 80 pixels); frame rate = 3.3 frames/s. The displayed area is 60 × 50 nm^2^. White double-headed arrows indicate the NTD dimers.

**Extended Data Fig. 4:**
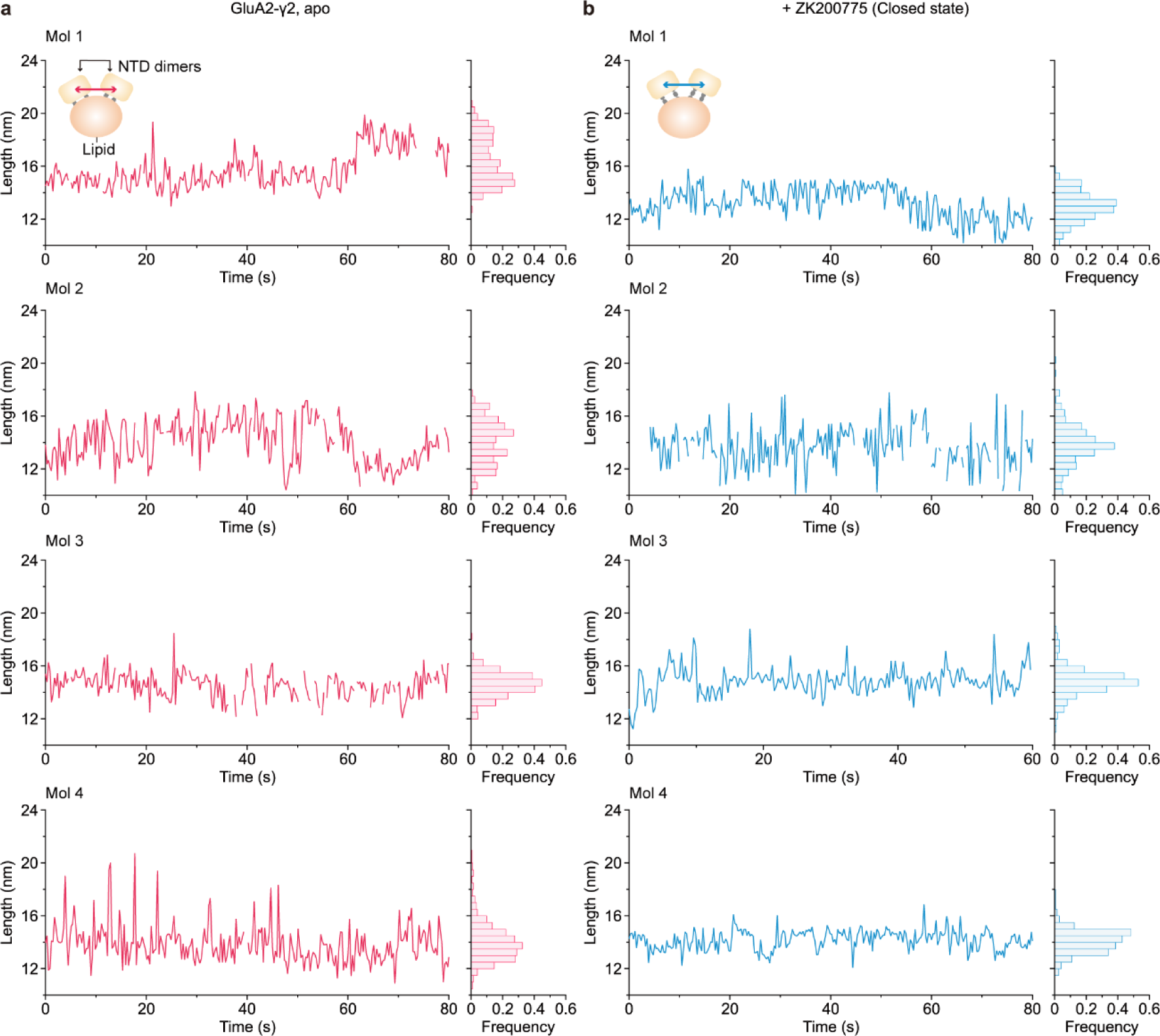
Fluctuation of the NTD dimers in GluA2-γ2 in the apo and with a competitive antagonist, related to Fig. 2. **a, b**, Four representative time courses of the changes in distance between NTD dimers in GluA2-γ2 apo (**a**) and ZK200775-bound (**b**) forms. Blank areas were not measurable due to NTD-dimer splitting.

**Extended Data Fig. 5:**
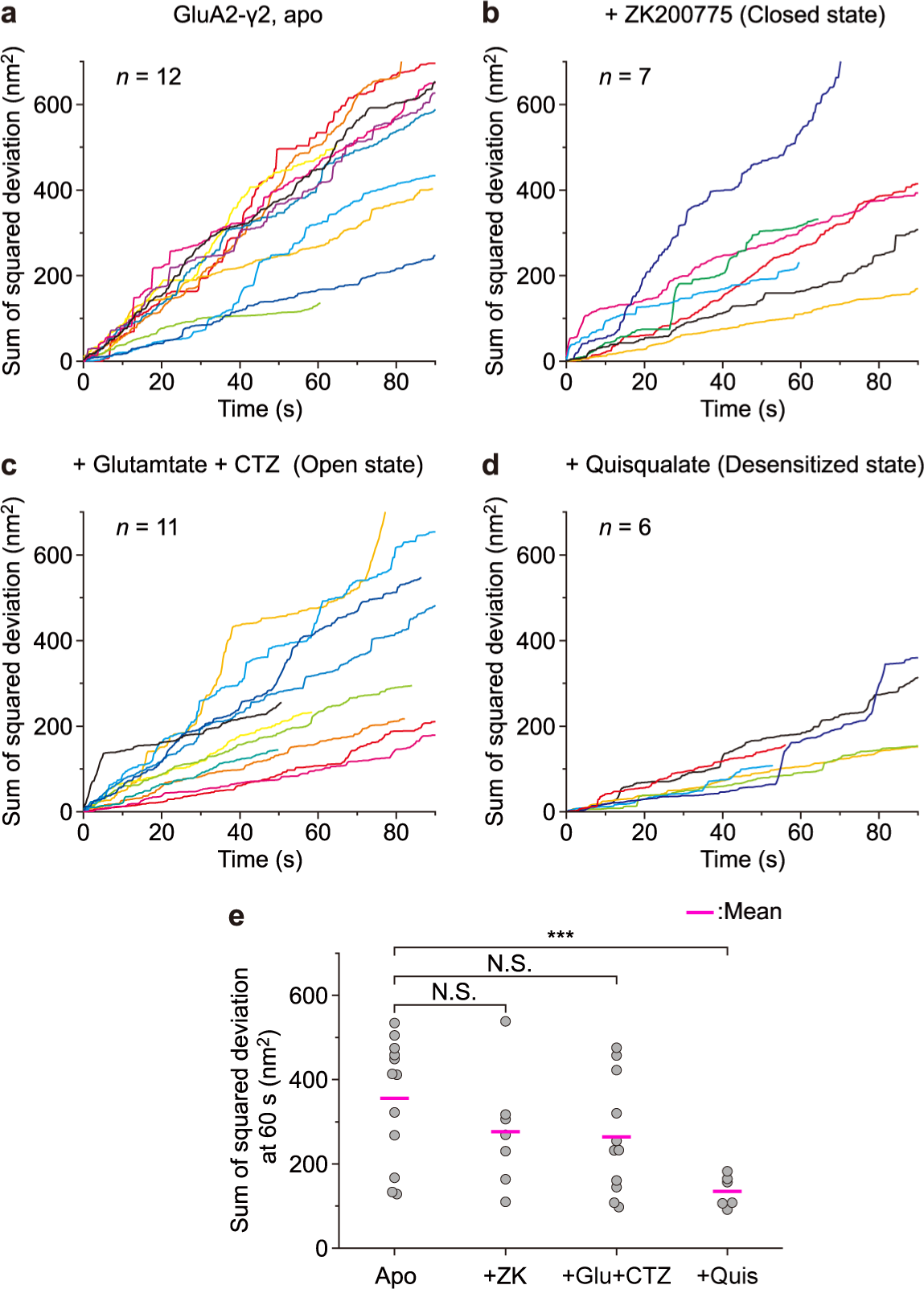
Fluctuation of the NTD dimers in GluA2-γ2 was restricted in the desensitized state, related to Fig. 2. **a-d**, Sum over time of squared deviation of the distance between NTD dimers in GluA2-γ2 apo (**a**), ZK200775-bound (**b**), Glu+CTZ-bound (**c**), and Quis-bound (**d**) forms. **e**, Sum of squared deviation of the distance between NTD dimers in GluA2-γ2 under different experimental conditions at appropriately 60 s; not significant (N.S.); \*\*\**P* < 0.001 (Kruskal‒Wallis test with Dunnett post hoc test).

**Extended Data Fig. 6:**
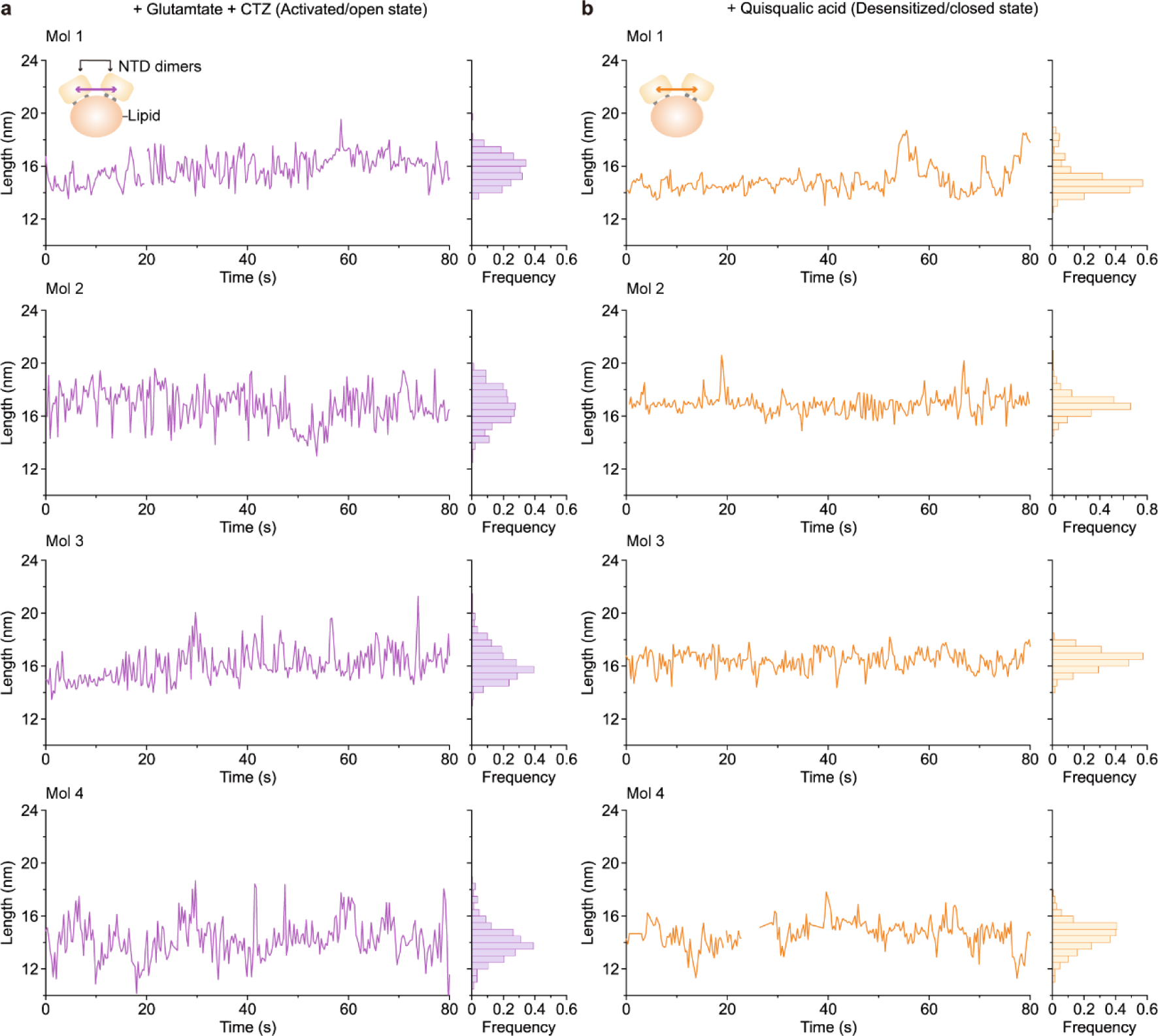
Fluctuation of the NTD dimers in GluA2-γ2 in the activated/open and desensitized/closed states, related to Fig. 2. **a, b**, Four representative time courses of changes in distance between NTD dimers in GluA2-γ2 bound with Glu and CTZ (**a**) and with Quis (**b**). Blank areas were not measurable due to NTD-dimer splitting.

**Extended Data Fig. 7:**
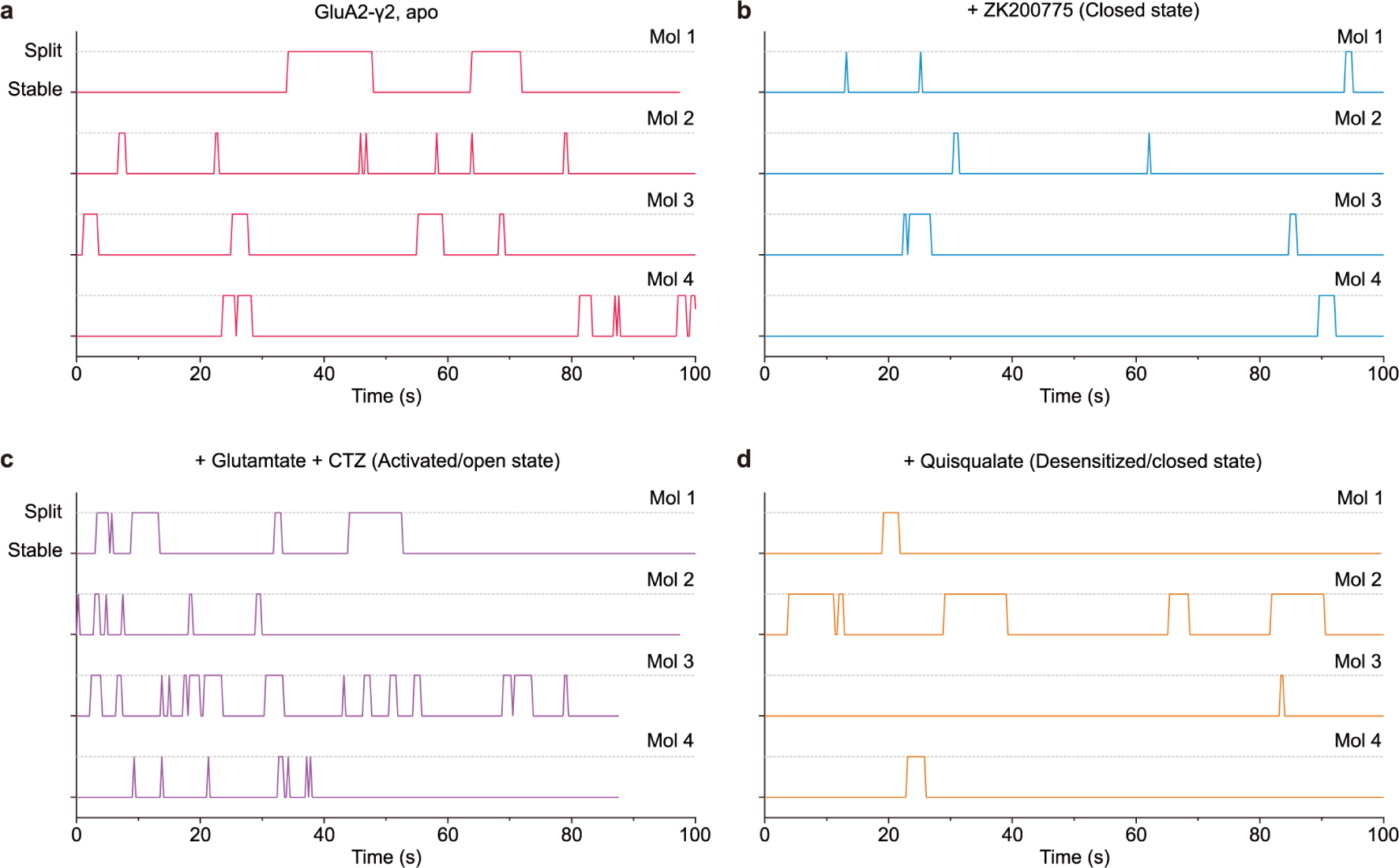
Kinetics of the NTD-dimer splitting of GluA2-γ2, related to Fig. 3. **a-d**, Four representative time courses of the transition of stable and split states of the NTD dimers under different experimental conditions. Dotted lines indicate the split state.

**Extended Data Fig. 8:**
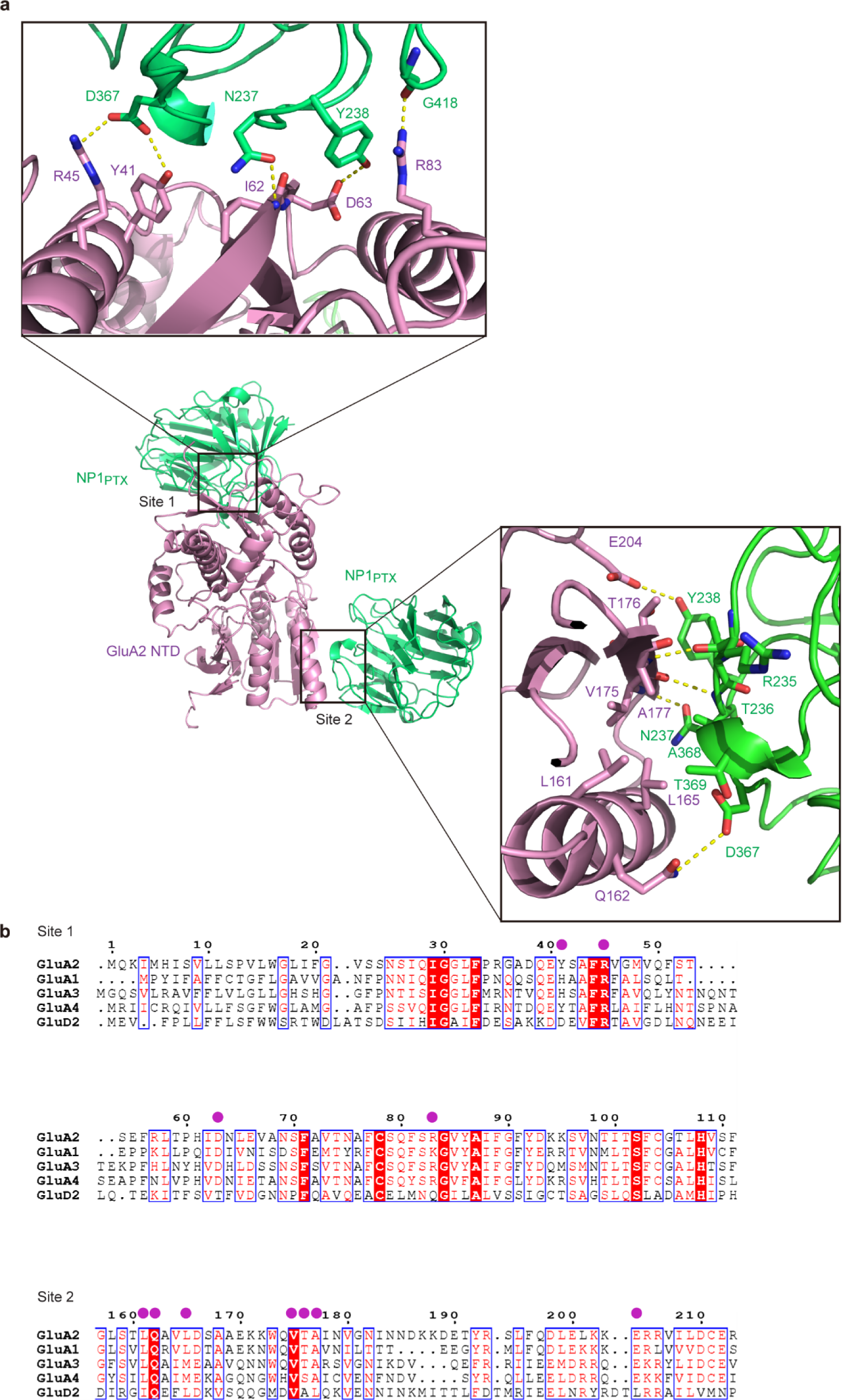
Binding interfaces between the GluA2 NTD and NP1_PTX_, related to Fig. 7. **a,** Close-up views of the binding interfaces between the GluA2 NTD and NP1_PTX_ as predicted by AlphaFold2. The overall structure and two potential interfaces (sites 1 and 2) are shown. Residues located at the interfaces are shown as sticks, and dotted lines indicate hydrogen bonds. **b,** Sequence alignment of rat GluA2 (P19491), GluA1 (P19490), GluA3 (P19492), GluA4 (P19493), and GluD2 (Q63226). Purple circles indicate residues predicted to be involved in the hydrogen bonds between the GluA2 NTD and NP1_PTX_.

**Extended Data Fig. 9:**
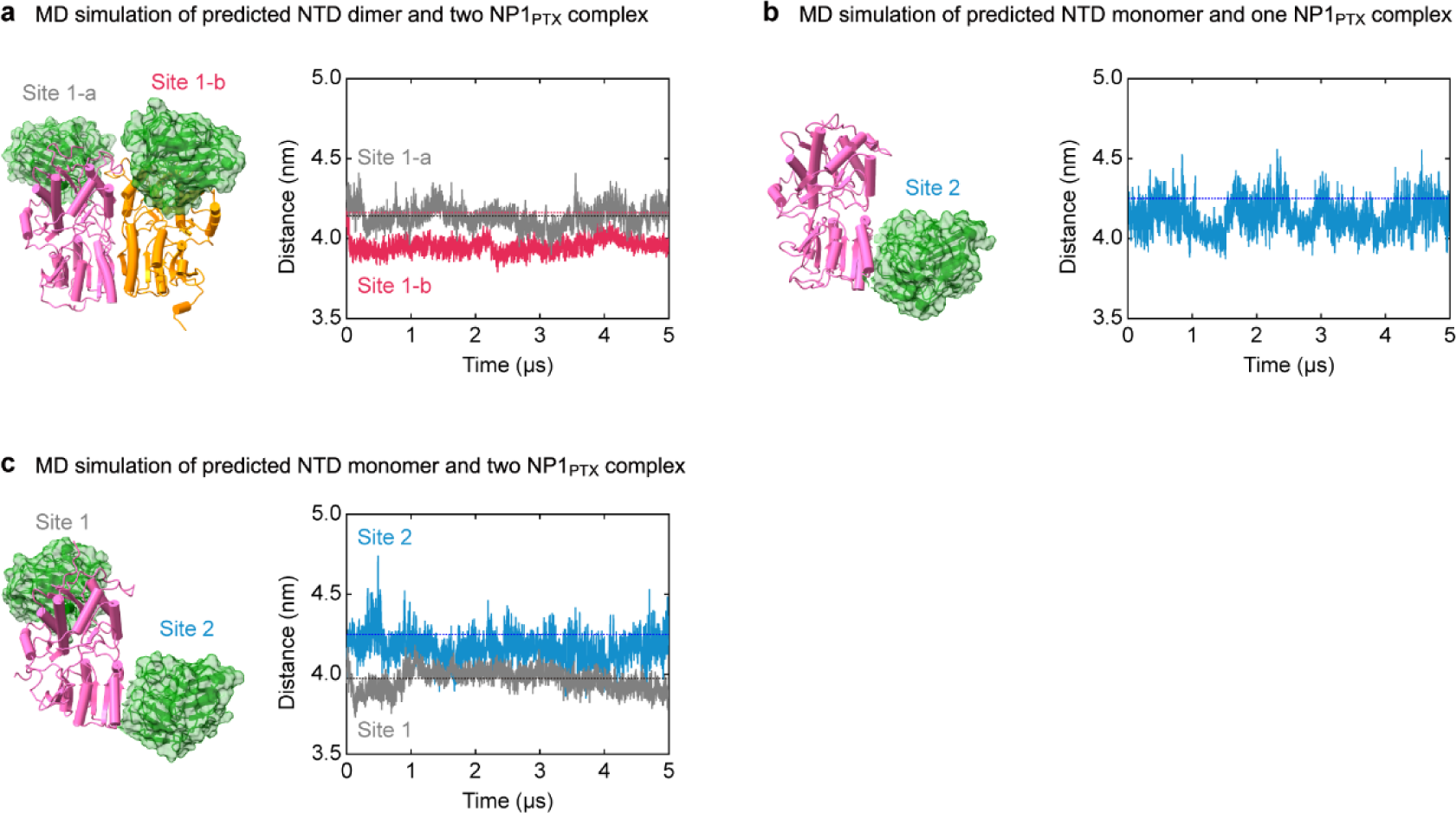
MD simulations of the NTD dimer or monomer and NP1_PTX_ complexes starting from the configurations predicted by AlphaFold2, related to Fig. 7. **a-c,** MD simulations of predicted NTD dimer–two NP1_PTX_ (**a**), NTD monomer–one NP1_PTX_ (**b**), and NTD monomer–two NP1_PTX_ (**c**). The time courses of the distance between the centre of all atoms in NP1_PTX_ and the centre of all atoms in the NTD monomer for each NP1_PTX_ are shown. In an NTD dimer, distance is defined as the shortest distance between the NTD monomer and NP1_PTX_ (**a**). The distances in the configuration predicted by AlphaFold2 are indicated by dashed lines.

**Extended Data Fig. 10:**
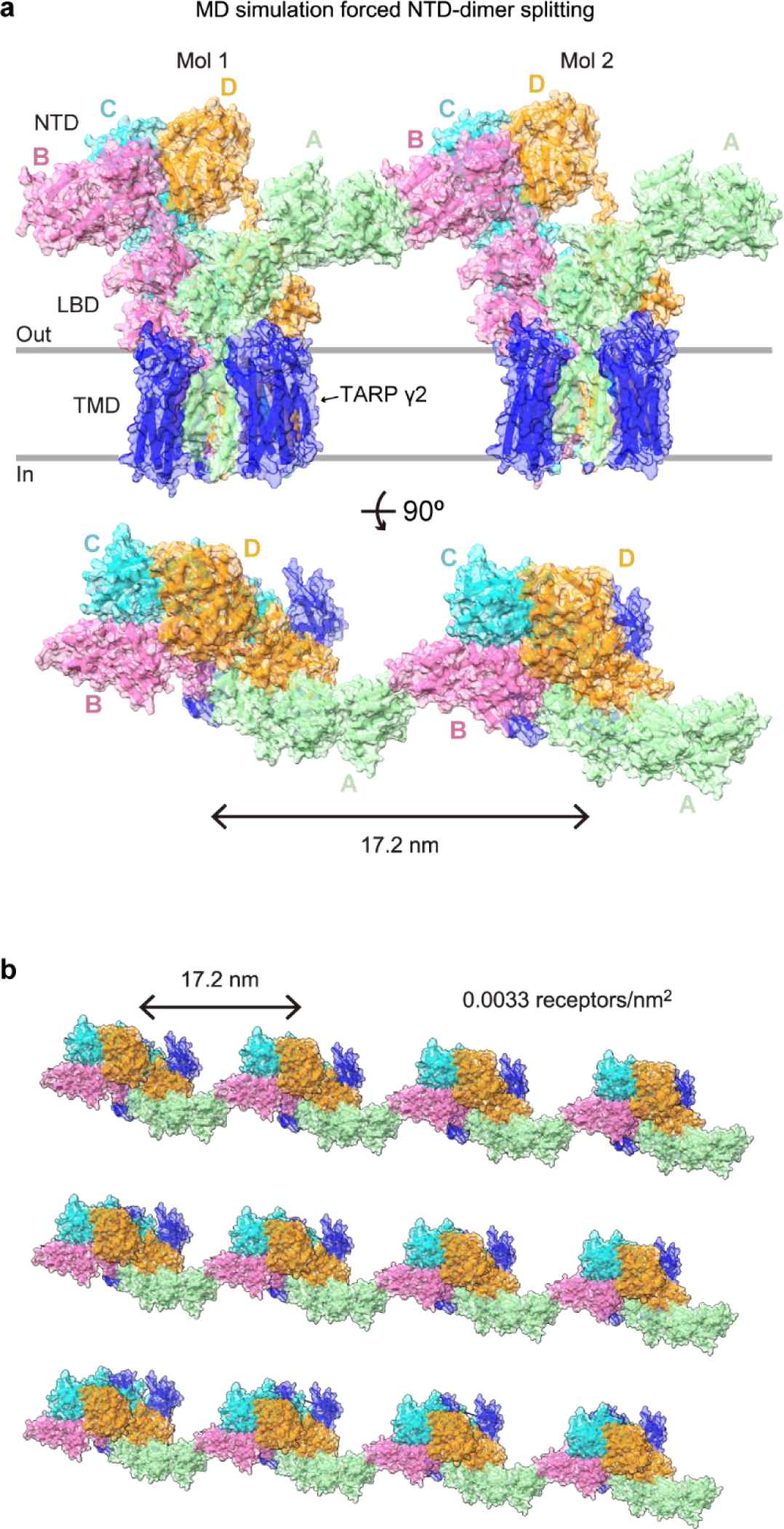
A configuration of interdimer exchange found in MD simulations after forced NTD-dimer splitting, related to Fig. 5. **a**, One NTD dimer (A-D) was forced to split. After splitting, the NTD monomer (A) in a GluA2-γ2 molecule (Mol 1) spontaneously interacted with the NTD monomer (B) in Mol 2 across the periodic boundary condition. Note that the density of GluA2-γ2 in the simulation area (0.0033 receptors/nm^2^) is similar to that in rat hippocampal neurons (0.0035 receptors/nm^2^). This interaction was maintained for approximately 200 ns. **b**, Top view of 3 x 4 GluA2-γ2 molecules aligned at 17.2 nm. At this density, neighbouring GluA2-γ2 molecules interact with each other through NTD-dimer splitting.

