## Supplemental Figs for "High-speed AFM reveals fluctuations and dimer splitting of the N-terminal domain of GluA2-γ2"

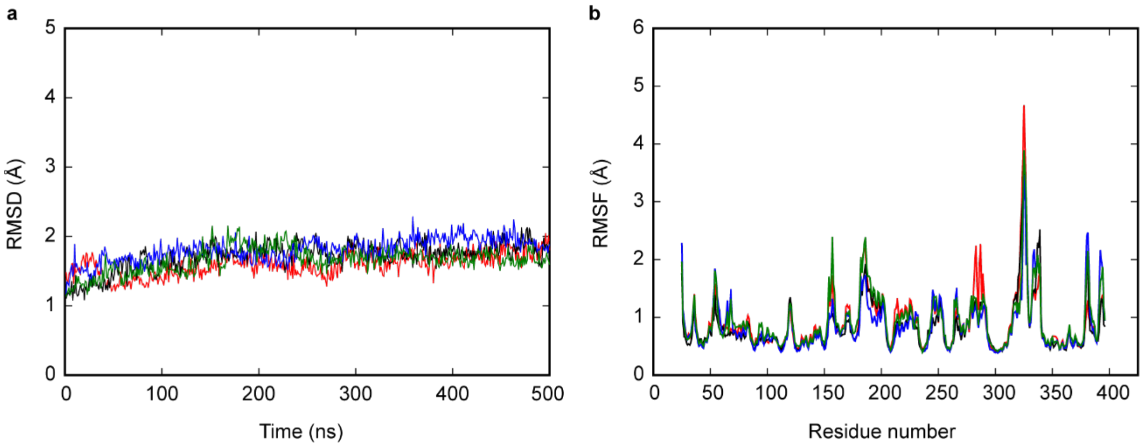


**Supplementary Fig. S1: The root mean square deviation (RMSD) and root mean square fluctuation (RMSF) of alpha carbon atoms in the NTDs, related to Fig. 5.**

**a**, RMSD from the crystallographic structure.

**b**, RMSF.

The RMSDs and RMSFs of the four NTD monomers are shown in different colours.

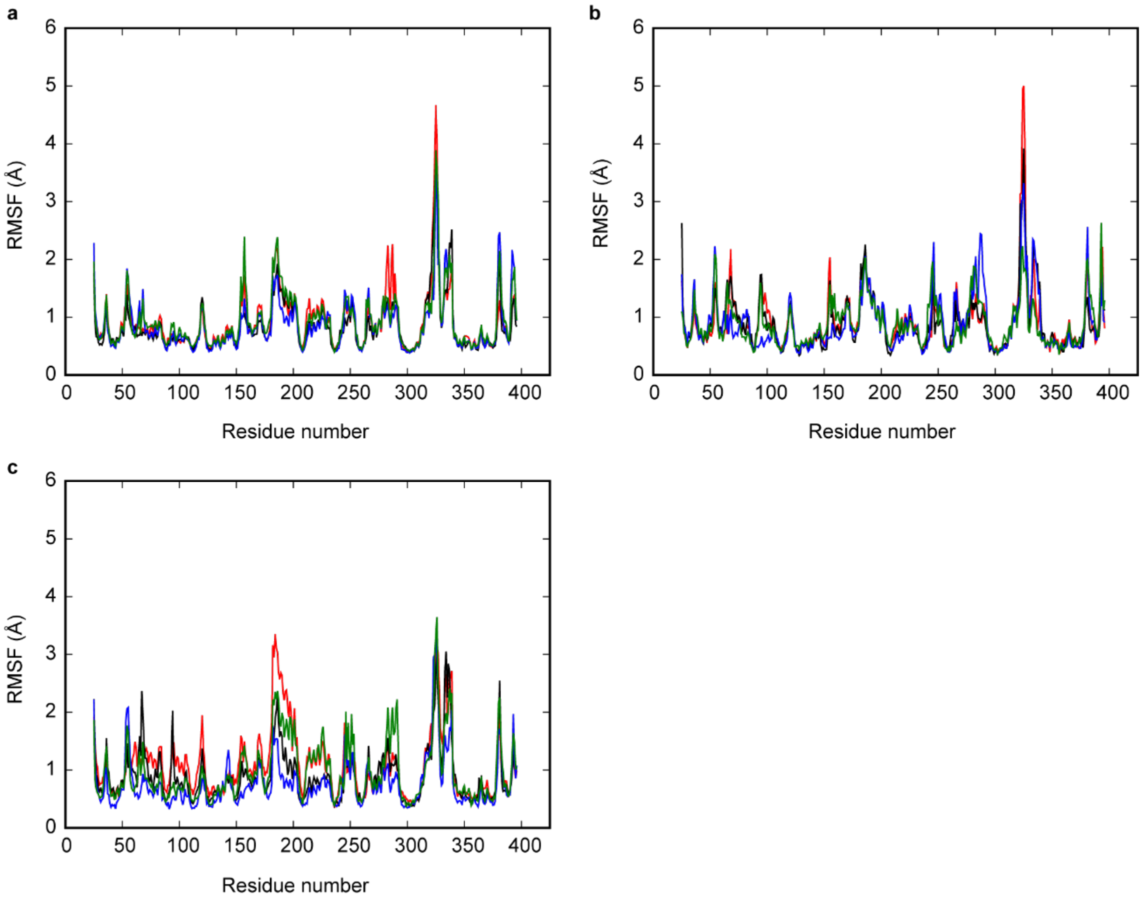


**Supplementary Fig. S2: The root mean square fluctuation (RMSF) of alpha carbon atoms in the NTDs after NTD-dimer splitting, related to Fig. 5.**

**a**, RMSF when RC = 3.6 nm, which is the same distance found in the crystallographic structure.

**b**, RMSF when RC = 7.0 nm.

**c**, RMSF when RC = 10.0 nm.

The RMSDs and RMSFs of the four NTD monomers are shown in different colours.

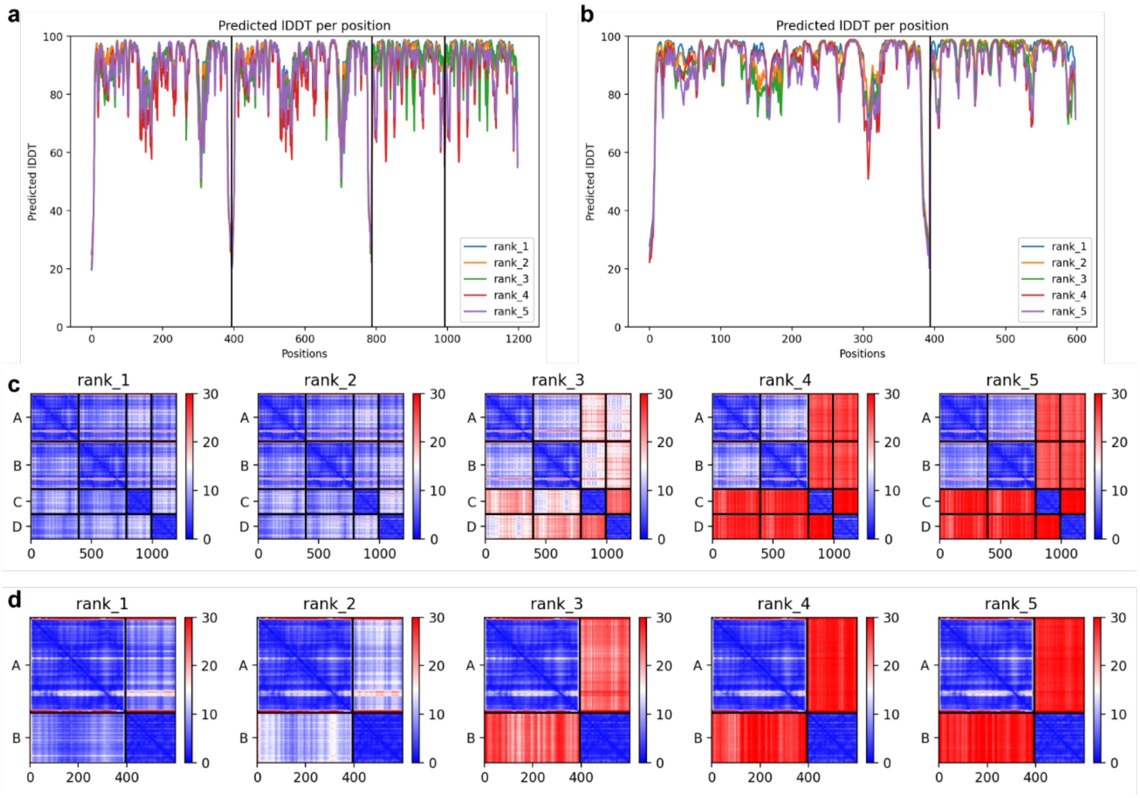


**Supplementary Fig. S3: AlphaFold2 prediction statistics, related to Fig. 7.**

**a, b**, Predicted local distance difference test (pLDDT) scores for each amino acid position of the GluA2 NTD dimer in complex with two $NP1_{PTX}$ molecules (**a**) and the GluA2 NTD monomer in complex with a single $NP1_{PTX}$ molecule (**b**).

**c, d**, Predicted aligned error maps for the GluA2 NTD dimer in complex with two $NP1_{PTX}$ molecules (**c**) and the GluA2 NTD monomer in complex with a single $NP1_{PTX}$ molecule (**d**).

**Supplementary Table S1: Parameters of the correlation function of NTD-dimer splitting, related to Fig. 4.**

$A_1\exp(-t/\tau_1) + A_2\exp(-t/\tau_2) + A_3\exp(-t/\tau_3)$

$\tau_{01}$ means the lifetime from stable to split states, while $\tau_{10}$ means the lifetime from split to stable states. Since the total time for most HS-AFM observations was approximately 100 s, the A3 components were negligible.

ZK, ZK200775; CTZ, cyclothiazide; Glu, glutamate; Quis, quisqualate.

| | $A_1$ | $A_2$ | $A_3$ | $\tau_1$ (s) | $\tau_2$ (s) | $\tau_3$ (s) | $A_1$ | | $A_2$ | | $A_3$ | |
|---|---|---|---|---|---|---|---|---|---|---|---|---|
| | | | | | | | $\tau_{01}$ (s) | $\tau_{10}$ (s) | $\tau_{01}$ (s) | $\tau_{10}$ (s) | $\tau_{01}$ (s) | $\tau_{10}$ (s) |
| apo | 0.450 | 0.497 | 0.069 | 0.91 | 4.3 | 186 | 25.8 | 0.94 | 121 | 4.4 | 5311 | 193 |
| +ZK (closed) | 0.251 | 0.750 | 0.014 | 1.2 | 50 | 84 | 18.2 | 1.3 | 749 | 53 | 1276 | 90 |
| +Glu +CTZ (open) | 0.187 | 0.277 | 0.536 | 0.46 | 4.7 | 183 | 6.1 | 0.50 | 61 | 5.0 | 2427 | 198 |
| +Quis (desensitized) | 0.132 | 0.839 | 0.026 | 0.53 | 6.6 | 735 | 16.4 | 0.55 | 205 | 6.8 | 22863 | 759 |

**Supplementary Table S2: Information on the MD simulation system of GluA2-γ2 and equilibrated box size, related to Fig. 5.**

| | |
|---|---|
| GluA2-γ2 | 1 |
| POPC | 887 |
| Water molecule | 161,547 |
| Glutamic acid | 4 |
| CTZ | 4 |
| $Na^+$ ion | 438 |
| $Cl^-$ ion | 438 |
| Total number of atoms | 666,719 |
| Ion concentration | 150 mM |
| Simulation box dimensions | 172.418 × 176.931 × 215.289 $Å^3$ |

**Supplementary Table S3: Information on the MD simulation systems of NTD-$NP1_{PTX}$ and equilibrated box size, related to Fig. 7.**

| | monomer | dimer | monomer + 2$NP1_{PTX}$ 2 |
|---|---|---|---|
| NTD | 1 | 2 | 1 |
| $NP1_{PTX}$ | 1 | 2 | 2 |
| Water molecules | 75,397 | 85,503 | 84,750 |
| $Na^{+}$ ion | 204 | 232 | 229 |
| $Cl^{-}$ ion | 196 | 216 | 217 |
| Total number of atoms | 235,926 | 275,735 | 267,234 |
| Ion concentration | 150 mM | 150 mM | 150 mM |
| Simulation box dimensions | 145.305 × 141.252 × 116.808 Å$^3$ | 151.898 × 139.805 × 132.419 Å$^3$ | 138.777 × 138.652 × 141.623 Å$^3$ |

### Supplementary video legends

**Video 1: HS-AFM videos of three representative apo-GluA2-γ2 molecules on mica, related to Fig. 1b.** GluA2-γ2 in the lipid without any addition of inhibitors or ligands. Magenta arrows indicate the splitting of the NTD dimers. Image size, 76 × 62 pixels$^2$; scan area, 60 × 50 nm$^2$; frame rate, 3.3 fps.

**Video 2: HS-AFM videos of three representative GluA2-γ2 molecules treated with ZK200775 on mica, related to Fig. 1c.** GluA2-γ2 in the lipid with 0.3 mM ZK200775. Image size, 76 × 62 pixels$^2$; scan area, 60 × 50 nm$^2$; frame rate, 3.3 fps.

**Video 3: HS-AFM videos of three representative GluA2-γ2 molecules treated with Glu and CTZ on mica, related to Fig. 2a.** GluA2-γ2 in the lipid with 3 mM Glu and 0.1 mM CTZ. Magenta arrows indicate the split of the NTD dimers. Green double-headed arrows indicate the NTD dimers after intradimer exchange. Image size, 76 × 62 pixels$^2$; scan area, 60 × 50 nm$^2$; frame rate, 3.3 fps.

**Video 4: HS-AFM videos of three representative GluA2-γ2 molecules treated with Quis on mica, related to Fig. 2b.** GluA2-γ2 in the lipid with 1 mM Quis. Magenta arrows indicate the splitting of the NTD dimers. Image size, 76 × 62 pixels$^2$; scan area, 60 × 50 nm$^2$; frame rate, 3.3 fps.

**Video 5: Movie of 500 ns MD simulation showing fluctuations of the NTDs and the LBDs in the bulk solution.** Each subunit of AMPAR is shown by a different colour. Glu and CTZ are shown by the space-filling model. Blue and purple balls are $Na^+$ and $Cl^-$ ions, respectively. Water molecules are shown as red dots and lipid molecules as tan line models. In the top view, only AMPAR is shown for clarity.

**Video 6: Movie of a 450 ns MD simulation showing fluctuations of the NTDs and the LBDs in the bulk solution after NTD-dimer splitting.** Each subunit of AMPAR is shown by a different colour. Glu and CTZ are shown by the space-filling model. Blue and purple balls are $Na^+$ and $Cl^-$ ions, respectively. Water molecules are shown as red dots and lipid molecules as tan line models. In the top view, only AMPAR is shown for clarity.

**Video 7: HS-AFM videos of two representative GluA2-γ2 in interdimer exchange of the NTD, related to Fig. 5b.** GluA2-γ2 in the lipid with 3 mM Glu and 0.1 mM CTZ. Magenta arrows indicate the splitting of NTD dimers. Green double-headed arrows and dotted boxes indicate the NTD dimers after interdimer exchange. Image size, 100 × 80 pixels$^2$; scan area, 80 × 64 nm$^2$; frame rate, 3.3 fps.

**Video 8: HS-AFM videos of four representative GluA2-γ2 and NP1 complexes with and without Glu and CTZ, related to Fig. 7d and e.** GluA2-γ2 in the lipid with 3 mM Glu and 0.1 mM CTZ. Magenta arrows indicate the beginning of the NTD-dimer split. Image size, 240 × 96 pixels$^2$; scan area, 120 × 96 nm$^2$ for the first three videos, 200 × 80 pixels$^2$; scan area, 80 × 64 nm$^2$ for the fourth video; frame rate, 3.3 fps.

**Video 9: Movies of 5 μs MD simulations showing interactions between the NTDs and NP1$_{PTX}$.** The NTD monomer or dimer is shown in purple, and NP1$_{PTX}$ is shown in green. (left) NTD dimer + two NP1$_{PTX}$, (middle) NTD monomer + one NP1$_{PTX}$, (right) NTD monomer + two NP1$_{PTX}$.

**Supplementary Data 1: Amber library file of CTZ (CTZ.lib).**
All the charges and atom types of CTZ are written in the electronic supplement file of the AMBER library file.

**Supplementary Data 2:** The highest-ranked structural model of the GluA2 NTD monomer in complex with a single $NP1_{PTX}$ molecule by AlphaFold2 (binding at site 2 in Fig. 7).

**Supplementary Data 3**: The highest-ranked structural model of the GluA2 NTD dimer in complex with two $NP1_{PTX}$ molecules by AlphaFold2 (binding at site 1 in Fig. 7).

**Supplementary Data 4**: The distances between NTD and $NP1_{PTX}$ in 5 μs MD simulations.
